# Neurogenetic evidence that Dilp8 promotes developmental-stability via Lgr3-neuron oscillation

**DOI:** 10.1101/2024.11.26.625404

**Authors:** Rebeca Zanini, Joana Pereirinha, Magdalena Fernandez-Acosta, Andreia P. Casimiro, Mariana Pinho, Lara Lage, Andres Garelli, Fabiana Herédia, Alisson M. Gontijo

## Abstract

The ability to achieve a species-specific size and proportion despite developmental or environmental perturbations is termed developmental stability. The molecular and cellular processes behind this are best understood in insects. In *Drosophila,* a peripheral-tissue stress signal, the relaxin/insulin-like peptide Dilp8, promotes developmental stability during larval development via its neuronal receptor, Lgr3, an ortholog of vertebrate relaxin receptors. Lgr3 signaling is widely accepted to occur in–and to activate (depolarize)–the central brain growth-coordinating interneurons (PIL/GCL neurons). Here, using neurogenetic approaches, we confirm the requirement of Lgr3 in PIL/GCL neurons, but unexpectedly find that they require both silenced (hyperpolarized) and active (depolarized) states for an appropriate response to Dilp8. These results are most simply explained if Lgr3 activation by Dilp8 triggers PIL/GCL-neuron oscillatory activity, and such oscillations promote developmental stability. PIL/GCL neurons express and require Cyclin A–which can form cell-cycle oscillator complexes with cyclin-dependent kinases–for their response to Dilp8, independently of Rca1 (regulator of CycA)/Emi1 (early mitotic inhibitor). This opens the possibility that cell-cycle machinery can be co-opted for postmitotic neuron oscillations, adding to an increasing list of postmitotic roles for cyclins. Neuroanatomically, we show that PIL/GCL neurons form reciprocally-innervating loops, which are common architectures in oscillating circuits and central pattern generators. The role of PIL/GCL neurons in developmental stability mirrors other homeostasis-regulating, peptide-driven oscillatory circuits found in the vertebrate hypothalamus, a developmentally-homologous region to the one occupied by PIL/GCL neurons in the fly brain.

## INTRODUCTION

Animals coordinate the growth of their body and body parts using both local and systemic programs to reach a species-typical final size and proportion [1-5]. The robustness of this process against intrinsic or environmental perturbations is termed developmental stability [1,2,6,7]. In *Drosophila*, the relaxin/insulin-like peptide 8 (Dilp8) mediates interorgan growth coordination during larval development [8-11]. Dilp8 is produced and secreted from larval peripheral epithelial tissues called imaginal discs (larval precursors of adult appendages such as eyes, legs, and wings) undergoing growth stresses, such as during tumorigenesis or regeneration. Dilp8 activity promotes developmental stability by delaying the onset of metamorphosis, which terminates disc growth. This delay provides extra time for the discs to compensate for their abnormal growth [12] and in its absence, animals accumulate developmental errors and/or die [8,10,13-16].

Dilp8 activity depends on the presence of the relaxin receptor-like protein, Leucine-rich repeat-containing G protein-coupled receptor (Lgr3), an orthologue of the vertebrate relaxin receptors RXFP1/2 [10,17,18], in larval neurons [19-22]. Two bilateral brain interneurons, named *pars intercerebralis Lgr3*-positive (PIL)/growth coordinating *Lgr3* (GCL) neurons, hereafter referred to as PIL/GCL neurons, have been hypothesized—but not functionally tested—to be the critical neurons mediating the Lgr3-dependent response to the Dilp8 peripheral tissue stress signal [10,19-22] (**Figure 1A**). This hypothesis is based on the following main pieces of evidence: 1) the Dilp8-dependent delay in the onset of pupariation can be suppressed by RNAi-mediated knock-down of *Lgr3* using GAL4-dependent gene drivers that drive expression in subpopulations (hundreds) of neurons that include the PIL/GCL neurons. Namely, these are the *GMR19B09-GAL4* (*R19B09>*) driver (*R19B09* is a ∼3.7-kb *cis*-regulatory motif (CRM) corresponding to the 7th intron and part of the flanking exons of the *Lgr3* locus [19-22,23]) (**Figure 1B**), *Lgr3-GAL4::VP16* (a *GAL4::VP16* insertion into a BAC containing the whole *Lgr3* gene plus ∼1-kb upstream and 6.6-kb downstream sequence, substituting the first exon of *Lgr3* and carrying its own new 3’UTR [19]), and *MZ699-GAL4* (*MZ699>*, an unmapped *GAL4* enhancer trap [20,24]). *MZ699>* and *R19B09>* expression overlaps in at least 9 bilateral cells in the central nervous system (CNS) [20], whereas an intersection between *R19B09>* and *Lgr3-GAL4::VP16* is reported to overlap only in PIL/GCL neurons [19]. The latter might be a matter of assay sensitivity as *Lgr3-GAL4::VP16* supposedly recapitulates expression from the whole *Lgr3* locus, which should include some or all of the *R19B09>* pattern that by itself drives detectable expression in hundreds of neurons [20]. Importantly, neither intersection was yet used to manipulate *Lgr3* activity and therefore test the hypothesis that *Lgr3* is required in the intersecting cells, and if so, in which of these cells. 2) Silencing of *Lgr3* in a substantial amount of ventral nerve cord (VNC) neurons using the *teashirt* (*tsh*)*-GAL4* driver, which drives expression in most VNC neurons with the notable exception of those in the segments A8/9, does not rescue the Dilp8-developmental delay caused by constitutive expression of *dilp8* under the control of a *tubulin* promoter (*Tub-dilp8*), suggesting that the critical cells that require Lgr3 to convey the Dilp8 response are located in the brain (such as the PIL/GCL neurons) or in segments A8/9 [20]. 3) Ubiquitous expression of Dilp8 activates gene expression from a cAMP (and Ca^2+^)-responsive element (CRE) reporter in PIL/GCL neurons [20,21], and this activation is dependent on neuronal *Lgr3* [21], as panneuronal RNAi against *Lgr3* abrogates CRE reporter activity in PIL/GCL neurons. Critically, whether or not this cAMP/Ca^2+^-dependent response is important or not for the Dilp8-Lgr3 developmental-stability pathway has not been investigated. 4) Neuronal silencing of *R19B09>-* or *MZ699>-*positive cells by driving the human *inwardly-rectifying potassium channel KCNJ2* (*Kir2.1*) [25,26] partially suppresses the tissue-damage-induced delay in the onset of metamorphosis [20], whereas the expression of the *Bacillus halodurans sodium channel NaChBac* under the control of *R19B09>* (which should cause electrical hyperexcitation of these neurons) caused a delay in the onset of metamorphosis [21].

**Figure 1.**
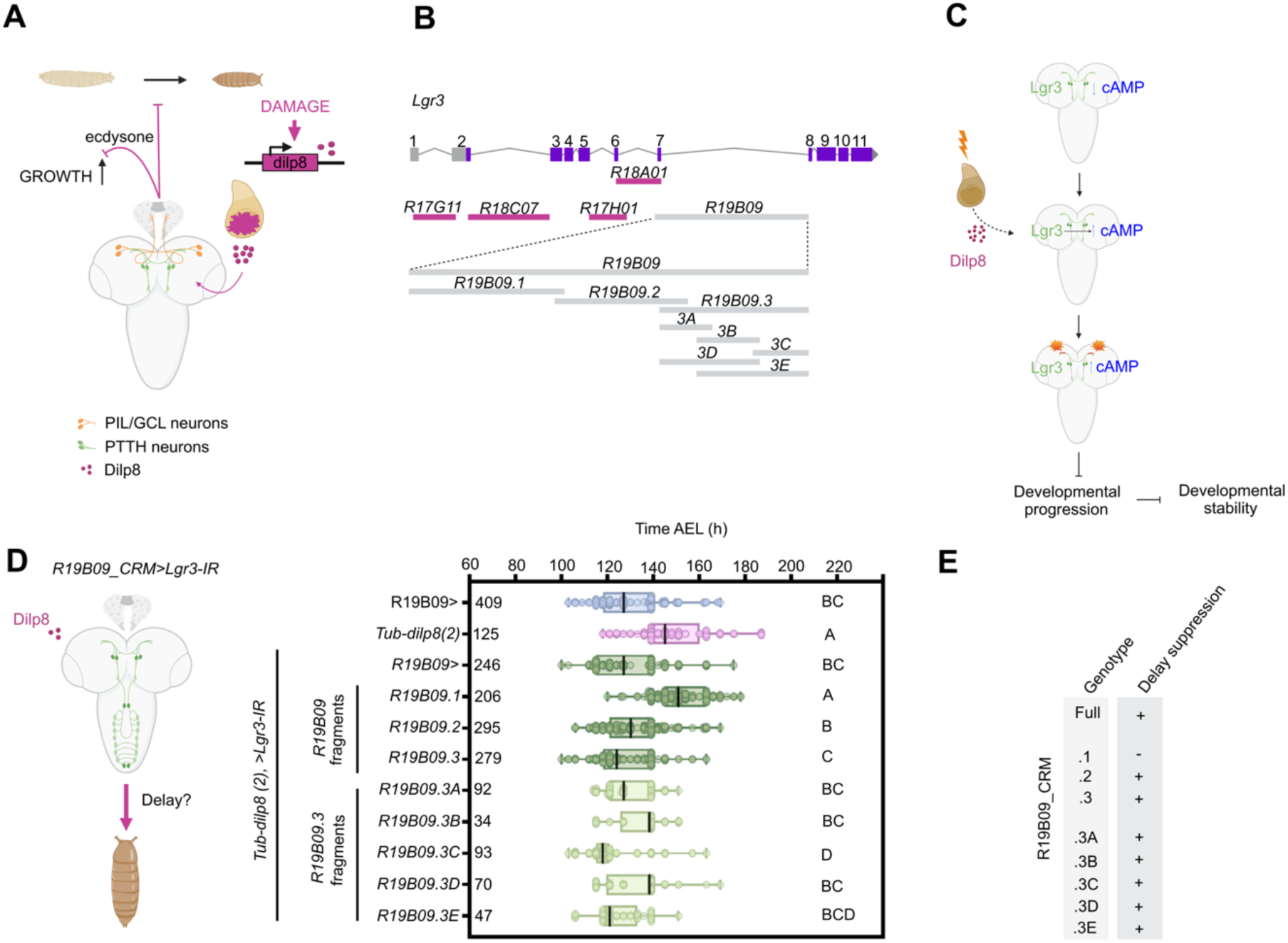
Enhancer bashing identifies specific fragments driving expression in cells requiring Lgr3 for the Dilp8-dependent developmental delay. (A) Model of the Dilp8-Lgr3 pathway for developmental stability in *Drosophila*. Scheme adapted from [10]. Dilp8 secreted from abnormally-growing imaginal discs indirectly inhibits ecdysone production, whose lower levels promotes developmental stability by delaying the transition from larval to pupal stage, allowing more time for imaginal discs to fine-tune their growth. (B) Schematic representation of the *Lgr3* gene depicting the *Lgr3* CRMs (*R17G11*, *R18C07*, *R17H01*, *R18A01*, and *R19B09*), and the bashed *R19B09* fragments (*.1-.3A-E*). (C) The current model of Lgr3-dependent activity in PIL/GCL neurons suggests that these are in an inactive state (inactive zone, blue) with low concentrations of intracellular cAMP, until Lgr3 is activated by the Dilp8 peripheral tissue stress signal. This activation leads to an increase of cAMP followed by membrane depolarization and synaptic activation (orange star represents synaptic activation; active zone, orange). In this model, PIL/GCL neuron activation induces a delay in developmental progression (larval to pupal transition), which promotes developmental stability. (D) Functional dissection of the *R19B09* CRMs identifies two independent regions that redundantly control *Lgr3* expression in cells that are relevant for the Dilp8-dependent delay. Box plot showing pupariation time (Time after egg laying (AEL)) in h of (N) larvae expressing *Tub-dilp8(2)* and *Lgr3-IR* under control of the *Lgr3* CRM, *R19B09* (*R19B09>*, control for suppression), and various *R19B09_CRM*s. Negative control (blue): *R19B09>* crossed with *w[1118]*. Positive control for the delay (magenta): *Tub-dilp8(2), UAS-Lgr3-IR* crossed with *w[1118]*. Whiskers extend to minimum or maximum values. Each dot represents one individual. The vertical bars represent the median. *P* < 0.0001, One-way ANOVA test. Genotypes sharing the same letter are not statistically different at α = 0.05, Tukey’s *post-hoc* test. (E) Summary of the phenotype of *Tub-dilp8* suppression for each *R19B09_CRM*. In the table, “+” denotes a positive finding, and “-” denotes a negative finding.

Hence, the generally accepted model is that Lgr3 receptor activation in the PIL/GCL neurons leads both to an increase in cAMP signaling and to PIL/GCL neuron activation by membrane depolarization and supposedly release of neurotransmitters to downstream neurons (**Figure 1C**). Activated PIL/GCL neurons then send inhibitory signals, probably via other neurons, such as PTTH neurons [19-21], which antagonize the biosynthesis of ecdysone by the ring gland, causing a delay in the onset of metamorphosis (**Figure 1A**).

As mentioned above, despite these findings and model, there are no reports describing the manipulation of *Lgr3* activity exclusively in PIL/GCL neurons to determine if these are indeed the neurons responsible for the developmental-delay activity of the Dilp8-Lgr3 pathway. Alternative models are that Lgr3 is required in other neurons or in other neurons in addition to the PIL/GCL neurons. Similarly, cAMP activity has not yet been functionally implicated in the Dilp8 delay in these neurons, as an alternative interpretation of the results described above would be that the Dilp8- and Lgr3-dependent CRE activation observed previously could be an artifact of the tools used and/or a true biological response to Dilp8-Lgr3 pathway activation, which is nevertheless unnecessary for the Dilp8-Lgr3 pathway function on developmental stability *per se*. Finally, the activity of the PIL/GCL neurons themselves has not been separately and exclusively genetically manipulated to test the model. Therefore, the question of what occurs downstream of Lgr3 activation at the neuronal-activity level remains open. Without direct tools to exclusively manipulate these neurons, the current model (**Figure 1C**) could be partially or completely incorrect, as the previously reported neuronal activity manipulations involved populations of hundreds of neurons.

Here, we used functional enhancer bashing and intersectional genetic experiments coupled with RNA interference (RNAi) and neurogenetic manipulations to test several aspects of this model. We generate PIL/GCL neuron genetic drivers with increased specificity and provide definitive functional evidence that downregulation of *Lgr3* in PIL/GCL neurons is sufficient to abrogate the developmental-delay activity of Dilp8, confirming the first part of the model. We also provide the first genetic evidence that a cAMP increase in PIL/GCL neurons via stimulatory G protein (G_s_) signaling is required for this Dilp8 activity, which is also consistent with the current model. However, in contrast to the current model, where Dilp8-triggered PIL/GCL-neuron depolarization was thought to delay the larval-to-pupal transition, we find that both hyper- and depolarized states are required for the developmental delay-promoting activity of these neurons in response to Dilp8. The simplest interpretation of these findings posits a model where Dilp8 triggers oscillatory behavior of PIL/GCL neurons in an Lgr3- and G_s_-dependent-manner, and that PIL/GCL membrane potential oscillation is critical to delay the larval-to-pupal transition and promote developmental stability. Immunohistochemical, genetic, and neuroanatomical studies provide an initial indication that the oscillation mechanism might involve the cell cycle-protein Cyclin A (CycA)–which has been previously linked to sleep length coordination in postmitotic neurons of adult flies [27-28]–and/or reciprocal loop network properties, widely known to mediate rhythmically-active neuronal networks (reviewed in [29-30]), respectively. Our results, apart from highlighting the importance of cell-type specific neurogenetic manipulations, reveal an unexpectedly complex role for PIL/GCL neurons in the Dilp8-dependent developmental stability pathway, provide new mechanistic insights into it, and point to an involvement of the Dilp8-Lgr3 relaxin-like pathway in the control of rhythmically active neuronal networks.

## RESULTS AND DISCUSSION

### DISSECTION OF THE R19B09 ENHANCER

#### R19B09 enhancer bashing identifies at least two independent cis-regulatory motifs capable of driving gene expression in cells requiring Lgr3 for the Dilp8-dependent delay

To start testing the model where Lgr3 is required in PIL/GCL neurons for the developmental delay in response to the Dilp8 peripheral stress signal, we attempted to generate PIL/GCL neuron-specific drivers to specifically manipulate these neurons using independent strategies. The first strategy consisted of enhancer bashing. As described above, the *Lgr3 R19B09* CRM/enhancer drives expression in a subpopulation of neurons that definitely contains the critical cells requiring *Lgr3* function to convey the Dilp8-dependent delay [19-21], and also cells that produce a developmental delay when depolarized [21]. Of the ∼270 *R19B09*>-expressing neurons [20,31], four are the PIL/GCL neurons [19-21]. *R19B09* was previously bashed into smaller partially-overlapping segments (*R19B09.1, R19B09.2*, and *R19B09.3*–the latter being further bashed into 5 fragments .*3A-E*) each of which was placed upstream of GAL4 (**Figure 1B**), and their expression pattern was characterized in the adult CNS [32], but not in the larval CNS. Here, we used these constructs to try to pinpoint the *R19B09* segment(s) that drive(s) expression in larval neurons where Lgr3 is necessary to convey the Dilp8-dependent delay and/or segment(s) that drive(s) expression in PIL/GCL neurons. Our hypothesis was that these will be the same, as is expected in the model (**Figures 1A and C**). The alternative is that the PIL/GCL neurons might not be the critical cells in which Lgr3 is required for the Dilp8-dependent delay, so that the fragments we identify while answering both questions might differ.

To answer the first question of which *R19B09* region(s) drive(s) expression in cells where Lgr3 is necessary to convey the Dilp8-dependent delay, we expressed RNAi against *Lgr3* (*Lgr3-IR* [33]) under the control of the different *R19B09 CRM-GAL4* (*R19B09_CRM>*), and assayed for their ability to suppress the delay produced by constitutive expression of *dilp8* using *Tub-dilp8(2)* (an insertion on the 2nd chromosome [21]) (**Figure 1D-E**). Consistent with previous results [20], *Tub-dilp8(2)* induced an 18-h median delay relative to controls, and the *R19B09>* control driver significantly rescued this delay when driving *Lgr3-IR*. These results further corroborated previous results showing that the Lgr3 receptor is necessary in *R19B09>-*expressing cells to delay the onset of metamorphosis in response to Dilp8 [19-21]. All *R19B09_CRM>* lines, except *R19B09.1>*, showed significant suppression of the Dilp8-induced delay when driving *Lgr3-IR* (**Figure 1D-E**). As the *R19B09.3A* and *R19B09.3C* regions correspond to non-overlapping segments of the seventh intron of *Lgr3* (**Figure 1B**), our result suggested there are at least two independent fragments within the *R19B09* element driving gene expression in cells requiring Lgr3 for the Dilp8-dependent delay. *Drosophila Lgr3* has hence evolved at least two enhancers in its seventh intron to secure Lgr3 protein expression in the cells required for the Dilp8-dependent delay. Whether or not these fragments drive redundant gene expression in the same population of Lgr3-positive, Dilp8-sensitive cells or not, and whether or not these are PIL/GCL neurons, cannot be concluded from these experiments alone.

#### The R19B09 CRMs that independently drive expression in the cells required for the Dilp8-dependent delay are shadow enhancers for the PIL/GCL neurons

To answer the second question of which *R19B09* segment(s) drive(s) expression in PIL/GCL neurons, we expressed membrane-targeted *myristoylated* (*myr)::tdTomato* [34] under the control of the different *R19B09_CRM>* lines in animals carrying the endogenous *Lgr3* translational reporter, *super folder green fluorescent protein*::*Lgr3* (*sfGFP::Lgr3*), which labels all Lgr3-positive neurons, including the PIL neurons [20] (**Figure 2A-B**). We crossed the different *R19B09_CRM>* lines with a stock carrying *UAS-myr::tdTomato; sfGFP::Lgr3*, and then performed immunohistochemistry of the F1 larval CNS to analyze the colocalization between *sfGFP::Lgr3* and *myr::tdTomato*, detected with an antibody against green fluorescence protein (GFP) and the endogenous tdTomato fluorescence, respectively. Therefore, when the *R19B09_CRM>* drives expression in the PIL neurons, we expect to observe a colocalization of both reporters. Our results show clear colocalization between *sfGFP::Lgr3* and *myr::tdTomato* driven either by *R19B09.3>* or by its five subfragments (*R19B09.3A-E*) (**Figure 2A**). Interestingly, these results show that PIL/GCL neuron expression is regulated by two independent and redundant CRMs contained in the non-overlapping fragments *R19B9.3A* and *R19B09.3C*. Redundant CRMs can be called shadow enhancers [35]. *Drosophila Lgr3* has hence evolved at least two shadow enhancers in its seventh intron to secure Lgr3 protein expression in the PIL/GCL neurons. Shadow enhancers can confer precision and robustness to gene expression, albeit complete spatial redundancy is rare [36]. Consistent with this, *R19B9.3A>* and *R19B09.3C>* seem to drive different patterns aside from the coinciding PIL/GCL neuron expression (**Figure 2A**).

**Figure 2.**
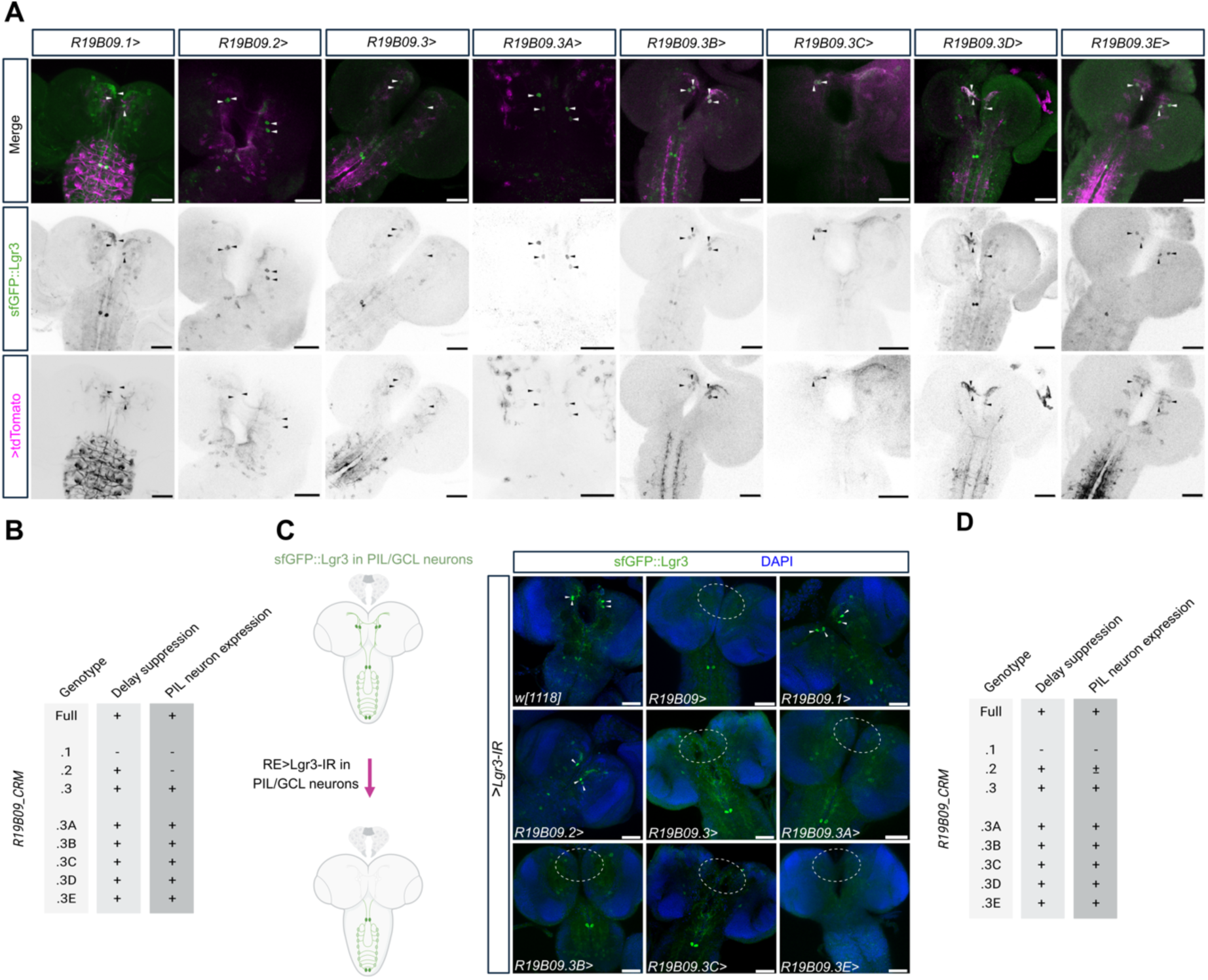
*Lgr3 R19B09.3 CRM*s drive expression in PIL neurons. **(A)** Sum of confocal z-stack slices stained with anti-GFP (green) to show *sfGFP::Lgr3* expression. Endogenous *myr::tdTomato* fluorescence driven by the various *R19B09_CRMs* is labeled in magenta. DAPI counterstain labels nuclei (blue). Both *R19B09.1>* and *R19B09.2>* do not drive detectable expression in sfGFP::Lgr3-positive PIL/GCL neurons, while the *R19B09.3>* fragments do. PIL/GCL neuron cell bodies are highlighted with arrowheads. **(B)** Summary of the phenotypes of *Tub-dilp8* suppression and PIL/GCL neuron expression for each *R19B09* CRM. In the table, “+” denotes a positive finding and “-‘’ denotes a negative finding. **(C)** Two independent CRMs within *R19B09* drive gene expression in the PIL/GCL neurons. Sum of confocal z-stack slices stained with anti-GFP (green) and DAPI (blue). The CRMs drive *Lgr3-IR,* reducing the *sfGFP::Lgr3* expression (green) in the subset of cells where they are expressed. As a negative control, we crossed *w[1118]* animals to *UAS-Lgr3-IR-V22, sfGFP::Lgr3* animals, so that *Lgr3-IR* is not expressed in this condition, and all sfGFP::Lgr3-positive neurons are labelled in green. As a positive control, we used *R19B09>* driving *Lgr3-IR*, expecting the absence of *sfGFP::Lgr3* in the PIL/GCL neurons. In the images where no PIL/GCL-like neurons are detectable, a white dashed circle denotes the anatomical region where their soma would otherwise be typically located. **(D)** Summary of the phenotypes of delay suppression and PIL neuron expression (summary of PIL neuron expression findings using both *sfGFP::Lgr3 Lgr3-IR* and *sfGFP::Lgr3 myr::tdTomato* colocalization techniques) for each R19B09 fragment. In the table “+” denotes a positive finding and “-‘’ denotes a negative finding. Scale bars, 50 µm.

Contrary to *R19B09.3>*, neither *R19B09.1>* nor *R19B09.2>* drove detectable expression of *myr::tdTomato* in the PIL/GCL neurons. As the *R19B09.1>* driver also does not suppress the Dilp8-dependent delay when driving *Lgr3-IR* (**Figure 1D-E**), we can also conclude that Lgr3 is not required in *R19B09.1>*-expressing cells, to convey the delay. However, in contrast to *R19B09.1>*, the *R19B09.2>* driver suppresses the Dilp8-dependent delay when driving *Lgr3-IR*. These initial results opened up the possibility that, contrary to the current model and to our expectations, the PIL/GCL neurons may not be the cells or may not be the only cells requiring Lgr3 in the Dilp8 pathway to induce a developmental timing delay.

Intrigued by the findings described above, we reasoned that our results could also be explained by the different sensitivity of the assays used above. Namely, that in the *Tub-dilp8-*delay experiments, certain *R19B09_CRM>* lines, such as the *R19B09.2>*, could drive very low levels of *Lgr3-IR* in the PIL/GCL neurons, which could nevertheless be sufficient to reduce Lgr3 protein levels enough to suppress the Dilp8-dependent delay, yet be too weak to lead to detectable myr::tdTomato fluorescence in the colocalization strategy. Hence, in order to independently confirm that *R19B09.1>* and *R19B09.2>* do not drive expression in the PIL/GCL neurons, and that the *R19B09.3>* element does, we devised a more sensitive immunofluorescence assay to test the capability of *R19B09_CRM>-*driven *Lgr3-IR* to reduce *sfGFP::Lgr3* protein levels in the PIL/GCL neurons. Namely, we crossed animals of the genotype *UAS-Lgr3-IR-V22, sfGFP::Lgr3* with the different *R19B09_CRM>* lines, and performed immunofluorescence analyses on dissected CNSs from the F1 larvae of the genotype *R19B09_CRM>Lgr3-IR; sfGFP::Lgr3*, using an antibody against GFP. Consistent with the results above using the myr::tdTomato reporter, virtually no detectable *sfGFP::Lgr3* was observed in cell bodies located in the PIL/GCL neuron anatomical region when *R19B09.3>* was used to express *Lgr3-IR* [0.08 ± 0.28, Mean ± standard deviation (SD) PIL/GCL neurons/brain; only 1 out of 13 brains had 1 PIL/GCL neuron detectable] (**Figure 2C-D**). Qualitatively similar results were found for all *R19B09.3>* subfragments tested, including *R19B09.3A>* and *R19B09.3C>* (**Figure 2C-D**). These results confirm that *R19B09.3* and both *R19B09.3A* and *R19B09.3C* shadow enhancers drive expression in the PIL/GCL neurons, and that this expression is sufficiently strong to reduce endogenous Lgr3 protein levels to undetectable levels when driving *Lgr3-IR*.

In contrast, driving *Lgr3-IR* under the control of *R19B09.1>* did not affect *sfGFP::Lgr3* expression in PIL/GCL neurons (3.90 ± 0.30 PIL/GCL neurons/brain, *N* = 11 brains), which demonstrates this segment does not drive significant GAL4 expression in PIL/GCL neurons, and is consistent with the lack of colocalization between *sfGFP::Lgr3* and *R19B09-1>*myr::tdTomato (**Figure 2A**). *Lgr3-IR* expression under the control of *R19B09.2*>, however, reduced *sfGFP::Lgr3* expression in PIL/GCL neurons, so that only a fraction of the four PIL/GCL neurons were detectable (2.16 ± 1.16 PIL/GCL neurons/brain, *N* = 6), suggesting that *R19B09.2>* drives weak, yet biologically-effective (capable of reducing endogenous Lgr3 protein levels) gene expression in PIL/GCL neurons. This very likely explains why *R19B09.2>Lgr3-IR* is sufficient to suppresses the Dilp8-dependent delay **(Figure 1C-D)**, and also provides an initial indication that the collective activity of the PIL/GCL neurons is critical for a proper response of Lgr3-positive neurons to the Dilp8 developmental-delay promoting signal. The fact that we could not see colocalization between R19B09.2>myr::tdTomato and sfGFP::Lgr3 **(Figure 2A-B)**, could be explained by a weaker sensitivity of the colocalization assay–*i.e.,* such that the expression of *myr::tdTomato* is too weak to be detected in our staining conditions. It is plausible that the weak PIL/GCL neuron expression conveyed by *R19B09.2* arises from its partial overlap with *R19B09.3A*, although we cannot exclude contributions from other upstream regulatory elements. Regardless, these results are highly consistent with the model that PIL/GCL neurons are the cells where Lgr3 is required to transduce the Dilp8-dependent delay.

#### R19B09 enhancer bashing identifies an incongruence between cells requiring Lgr3 and those inducing a developmental delay when depolarized

Previous work has shown that depolarization of *R19B09-*positive neurons using the NaChBac sodium channel was sufficient to delay the onset of metamorphosis [21]. If the PIL/GCL neurons are indeed the neurons causing such a delay, as hypothesized in the current model (**Figure 1A and C**), then the neurogenetic activation of all *R19B09_CRM>-*positive cells except *R19B09.1>* should cause a delay in development. To address this, we first attempted to independently replicate the finding above by thermogenetically activating *R19B09>-*positive neurons expressing the transient receptor potential cation channel A1 ortholog (TrpA1) with a temperature shift from 18°C to 29°C at the second larval instar stage **(Figure 3A)**. TrpA1 expression leads to cation influx in the cells and depolarization at 29°C [37-41]. To further test the epistatic relationship between neuronal activation and Lgr3 requirement, we performed these experiments in the presence or not of *Lgr3* RNAi. The results show that *R19B09>* significantly delays the onset of metamorphosis when driving TrpA1 **(Figure 3A)**, corroborating the previous report using an independent method of neuronal activation [21]. Furthermore, we find that the TrpA1-induced delay does not require Lgr3 in the *R19B09>*-expressing cells, because the delay is not rescued by co-expression of *Lgr3-IR* **(Figure 3A)**. This suggests that activation of R19B09 neurons is epistatic to Lgr3 activity in these neurons. One possibility is that firing of R19B09 neurons occurs downstream of Lgr3 activation by Dilp8, hence neuronal Lgr3 becomes dispensable. This is consistent with the current proposed model.

**Figure 3.**
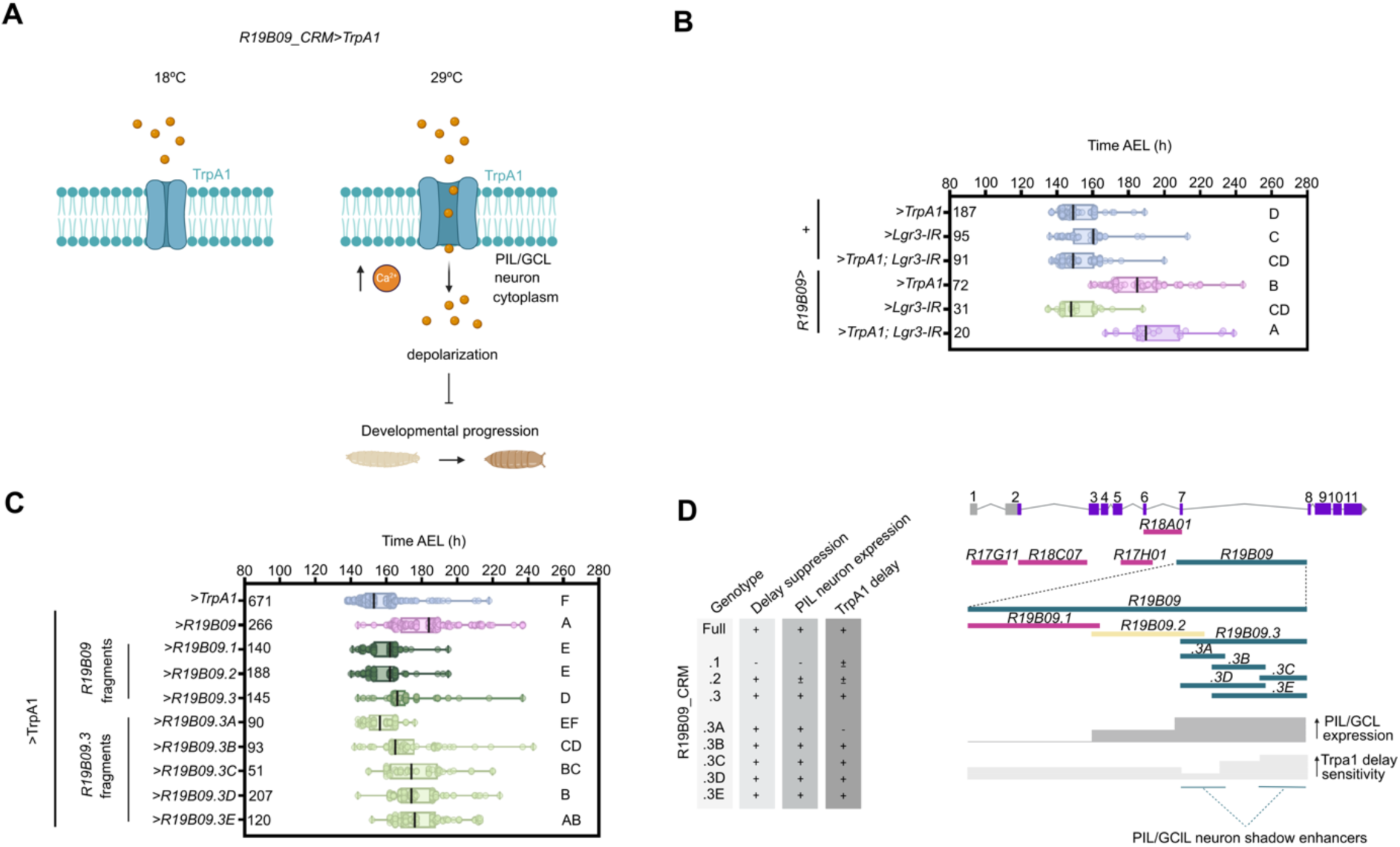
*Lgr3 R19B09.3* CRMs drive expression in TrpA1-sensitive cells, inducing a developmental timing delay. **(A)** Scheme depicting the experimental design for testing the sensitivity of the different cell populations marked by the *R19B09_CRMs* to the temperature-dependent Trpa1 neuronal activation. The final readout is timing of the larva to puparium transition (Developmental progression). Temperature-dependent TrpA1 activation (29 C) in PIL/GCL neurons will lead to cation influx (such as Calcium ions (orange circles) and neuronal depolarization, which is predicted to lead to a delay in the larval to pupariation transition according to the current model (Figures 1A **and C**). **(B)** Box plot showing pupariation time (Time after egg laying (AEL)) in h of (N) larvae expressing *TrpA1* and/or *Lgr3-IR* under the control of *R19B09>*. Negative controls (blue): each line crossed with *w[1118]*. Whiskers extend to minimum or maximum values. Each dot represents one individual. The vertical bars represent the median. *P* < 0.0001, One-way ANOVA test. Genotypes sharing the same letter are not statistically different at α = 0.05, Tukey’s *post-hoc* test. **(C)** Box plot showing pupariation time of larvae expressing *TrpA1* under the control of the various *Lgr3 R19B09_CRM* lines. Negative controls (blue): *UAS-TrpA1* crossed with *w[1118]*. Positive control (magenta): *R19B09>TrpA1*. *P* < 0.0001, One-way ANOVA test. Rest, as in (B). **(D)** Left: summary of the phenotypes of delay suppression, PIL/GCL neuron expression (compilation of PIL/GCL neuron expression findings using both sfGFP::Lgr3+Lgr3-IR and sfGFP::Lgr3+myr::tdTomato colocalization techniques) and TrpA1 delay for each *R19B09_CRM* fragment. In the table “+” denotes a positive finding and “-“ denotes a negative finding. Right: the *Lgr3* gene has at least two shadow enhancers that are active in the PIL/GCL neurons. The *R19B09* CRM was fragmented into three small elements: the *R19B09.1* fragment (magenta bar), which neither regulates the response to Dilp8 in Lgr3-positive neurons nor drives expression in PIL/GCL neurons; the *R19B09.2* fragment (yellow bar), which suppresses the Dilp8-dependent delay when driving *Lgr3-IR*, but its expression in PIL/GCL neurons is weak (PIL/GCL neuron expression, dark gray); and the *R19B09.3* fragment and its subfragments (dark cyan bars), which drive *Lgr3* in cells that regulate the Dilp8-dependent delay and drive expression in the PIL/GCL neurons, constituting at least two redundant “shadow” enhancers. The capability of the cells where these CRMs are active to cause a delay upon TrpA1 activation (TrpA1-sensitivity, light gray) increases more towards the 3’ region of the *R19B09.3* CRM with a dip around the *R19B09*.*3A* CRM region.

Next, we attempted to confirm if the PIL/GCL neurons are responsible for the R19B09>TrpA1-induced delay. Our strategy was to thermogenetically activate the different subsets of *R19B09>*-expressing cells using the different *R19B09_CRM>* lines crossed to *UAS-TrpA1* and score the timing of pupariation in similar temperature-shift conditions as the experiments described above **(Figure 3B-C)**. The results showed that all *R19B09_CRM>* lines, except *R19B09.3A>*, induced some extent of statistically significant delay in pupariation time when compared with the *UAS-TrpA1* negative control (**Figure 3B-C**). The weakest lines were *R19B09.1>* and *R19B09.2>* (∼9-h median delays relative to control, *P <* 0.05, Tukey’s *post-hoc* test for both). As all *R19B09.3*-derived lines, except for the *R10B09.3A>*, induced stronger delays upon TrpA1 activation (19.3 ± 4.3 h, average median delay ± SD), we interpret these results as indicative that the most relevant TrpA1-sensitive cells are part of the population of cells in which elements located at the two distal thirds of the *R19B09* element (represented by the CRM segments *R19B09.3, 3B, 3C, 3D,* and *3E*) are active. The fact that *R19B09.3A>-*expressing cells do not induce a delay when thermogenetically activated despite being clearly active in PIL/GCL neurons (**Figure 1B and Figure 2A-B**), posits two mutually exclusive scenarios. One is that thermogenetic activation of PIL/GCL neurons is sufficient to induce a delay, but it is blocked by depolarization of other neurons uniquely present in *R19B09.3A>*. In the alternative scenario, the thermogenetic activation of PIL/GCL neurons is not sufficient to induce a delay, and the experimentally extended larval period is the consequence of thermogenetic activation of other, yet unidentified cells, labeled by some *R19B09_CRM>* lines (e.g. *R19B09.3B>*) and absent in *R19B09.3A>*. To differentiate between both scenarios, we reasoned that if *R19B09.3A>* drove expression in cells that negatively affected developmental time, it should likewise negatively affect the delay caused by *R19B09>TrpA1*, but this was not the case (**Supplementary Figure S1**). Hence, the simplest explanation is that thermogenetic activation of PIL/GCL neurons is not sufficient to induce a delay. Future work could evaluate if the other populations of R19B09 neurons that do delay development when depolarized are part of the Dilp8- and Lgr3-dependent developmental stability pathway or rather represent cells acting in other pathways relevant for developmental timing control [3-4].

In summary, enhancer bashing of the *Lgr3 R19B09* CRM **(Figure 3D)** led to the four main conclusions: 1) that at least two PIL/GCL-neuron shadow enhancers (*R19B09.3A* and .*3C*) are present in the *Lgr3* gene; 2) there is a correlation between the *R19B09* CRMs that drive expression in cells requiring *Lgr3* for the Dilp8-dependent delay and those that drive expression in PIL/GCL neurons; 3) that a small *R19B09* region (*R19B09.3A*) that drives expression in PIL/GCL neurons is not sufficient to induce a delay upon thermogenetic activation; and 4) that other R19B09-positive cells, not marked by the driver *R19B09.3A>*, which excludes the PIL/GCL neurons, can delay the larval-to-pupal transition upon thermogenetic activation and in an *Lgr3-*independent manner.

### Targeting PIL/GCL neurons by intersectional genetics

#### A sparse gene driver intersection expressed in PIL/GCL neurons shows that removal of Lgr3 in PIL/GCL neurons is sufficient to abrogate its developmental delay activity

In the current model (**Figure 1A and C**), PIL/GCL neurons are strong candidates to mediate the Dilp8- and Lgr3-dependent developmental delay, but this had not been formally tested before this study. This is because PIL/GCL neurons had always been genetically manipulated together with hundreds of other cells due to the lack of PIL/GCL neuron-specific gene drivers. The enhancer bashing experiments described above are also consistent with this model, as they showed a correlation between the *R19B09* fragments that drive expression in cells requiring *Lgr3* for the Dilp8-dependent delay and those that drive expression in PIL/GCL neurons. Nevertheless, all of the *R19B09* fragments tested still drive expression in many other cells in addition to the PIL/GCL neurons. In order to independently confirm the model, we attempted to generate sparser and more specific PIL/GCL neuron drivers with intersectional genetics. Using fluorescent reporters, we had previously shown that the gene expression driven by the elements *R19B09-LexA* (*R19B09>>*) and *MZ699>* overlaps in at least 9 bilateral cells, two of which are the PIL/GCL neurons, while the other 7 neurons are in the VNC [20]. A genetic intersection between *R19B09>>* and *MZ699>* (*R19B09* ∩ *MZ699*) would thus allow us to functionally test if those 9 neurons are indeed required for the Dilp8-dependent delay by driving *Lgr3-IR* in the intersecting cells. To achieve this, we first quantified the genetic intersection of both drivers [*R19B09* ∩ *MZ699*] using a flip-OUT technique where *R19B09>>* drives *8xLexAop2-Flipase* (*FLPL*) [34] to remove the FRT-flanked transcriptional STOP sequence from the cassette *UAS-FRT-stop-FRT-GFP::myr* [42], to allow *MZ699>* to drive membrane-targeted GFP in the restricted population of cells in which both enhancer activities overlap. The *R19B09 ∩ MZ699>GFP* genetic intersection revealed an average of 16.5 ± 2.3 GFP-positive cells per hemi-CNS (average ± SD, *N* = 11 CNS preparations), not all of which were bilateral and/or consistently observed in all preparations. This number was considerably higher than the 9 bilateral neurons previously detectable by overlapping fluorescence derived from each driver activity independently [20], and could reflect the greater sensitivity of the genetic intersection method. Of these cells, 2.7 ± 0.7 (*N* = 22 hemibrains) were in the PIL/GCL neuroanatomical area (**Figure 4A**). These cells likely correspond to the two PIL/GCL neurons plus another “#5” neuron (MZ699 is active in all #5 neurons, of which the two PIL/GCL neurons are a subset [20]). Furthermore, we frequently observed another brain neuron (0.9 ± 0.6 per hemisphere), positioned more posteriorly to the PIL/GCL neurons (**Supplementary Figure S2**). In the VNC, an average of 9.2 ± 0.7 VNC neurons per side were positive and sometimes a neuroblast lineage was labeled (0.4 ± 0.3 per VNC side), typically in the lateral anteriormost VNC (**Supplementary Figure S2**). To further increase the specificity towards PIL/GCL neurons and allow genetic manipulations based on the intersection, we used another flip-OUT method based on the capability of GAL80 to directly inhibit GAL4 activity [43]. In this method, a *Tub-FRT-GAL80-FRT* flip-OUT cassette was used to convert *R19B09>>FLPL* expression into GAL4-permissive clones (GAL4 being driven by the *MZ699>* driver). In addition, to further restrict GAL4 activity to the brain, where the PIL/GCL neuron cell bodies are localized, we counteracted GAL4 expression in the VNC using *tsh*-*GAL80*, a GAL80 enhancer-trap insertion on the 5’UTR of *tsh* [44]. This intersection, “[*R19B09* ∩ *MZ699*] – *tsh>”*, is hereafter named *PIL/GCL-GAL4* (*PIL/GCL>*).

**Figure 4.**
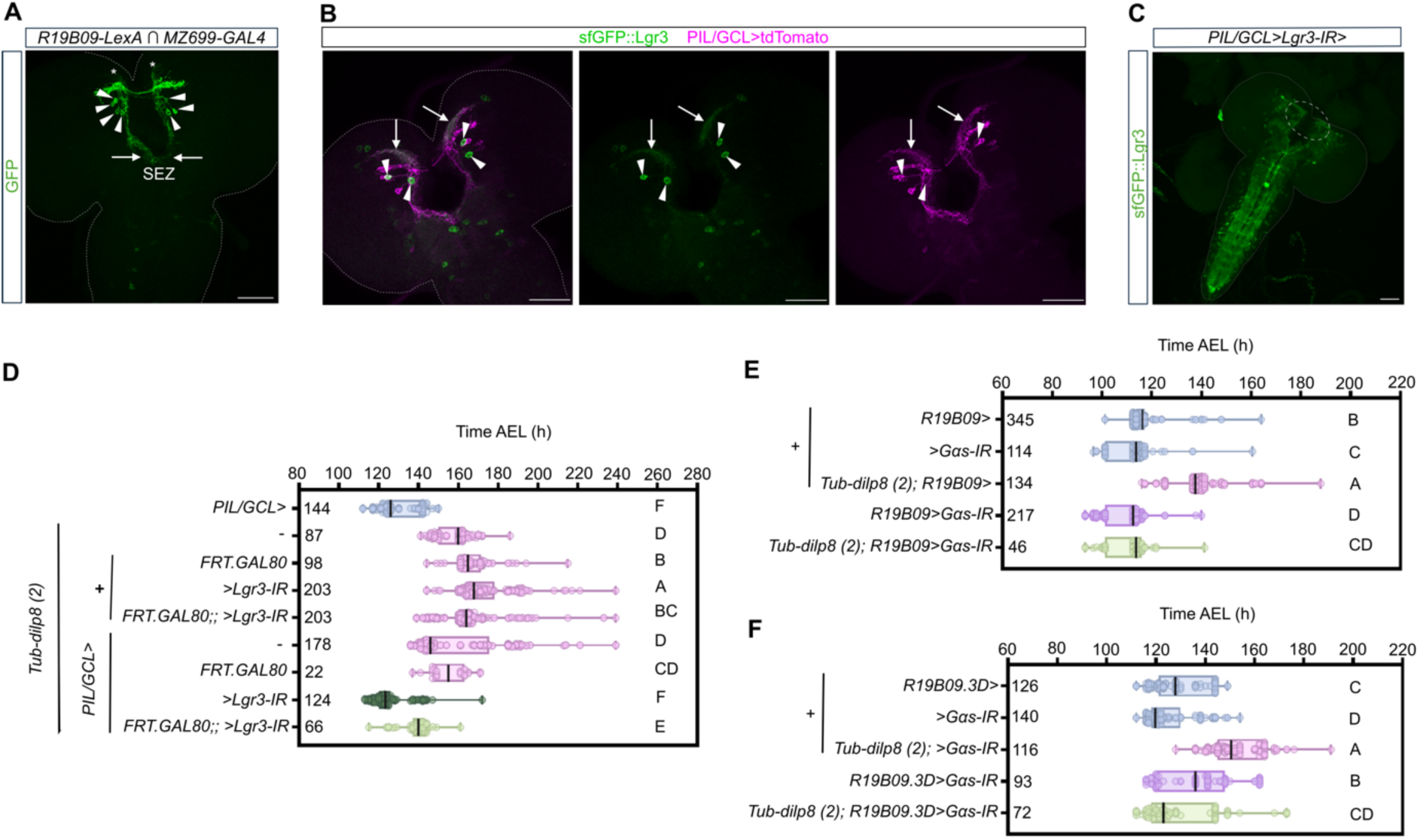
*Lgr3* and *Galphas* knockdown in PIL/GCL neurons is sufficient to abrogate its developmental delay activity. (**A**) The genetic intersection of *R19B09* and *MZ699* elements (*R19B09 ∩ MZ699*) reveals cells in the brain region corresponding to PIL/GCL neurons. Shown is the sum of confocal z-stack slices of an immunohistochemistry preparation of dissected wandering L3 larval CNSs (anterior contour depicted with a dashed white line) expressing *R19B09 ∩ MZ699>GFP::myr* and stained with anti-GFP (green). Arrowheads point to the soma of at least seven GFP-positive neurons, with contralateral anterior arborizations (asterisks), and ipsilateral descending projections (arrows) towards the subesophageal zone (SEZ). (**B**) PIL/GCL neurons are a subset of the #5 cells labeled with *PIL/GCL>.* Depicted is the sum of confocal z-stack slices of an immunohistochemistry preparation of dissected wandering L3 larval CNSs (anterior contour depicted with a dashed white line) stained with anti-GFP (green) to show the overlap between sfGFP::Lgr3 expression and PIL/GCL>myr::tdTomato (magenta). The contralateral projections of PIL/GCL neurons occupy the anteriormost and more curvilinear region of this neuropil compartment (arborizations with green and magenta overlap, arrows). The other straighter and less arborized contralateral projections correspond to those of other neighboring sfGFP::Lgr3-negative #5 group neurons (magenta). **(C)** *Lgr3* RNAi expression under the control of *PIL/GCL>* (*PIL/GCL>Lgr3-IR*) in the presence of the *sfGFP::Lgr3* cassette leads to a complete loss of detectable anti-GFP staining in the region corresponding to PIL/GCL neurons. Shown is the sum of confocal z-stack slices of an immunohistochemistry preparation of dissected wandering L3 larval CNSs (anterior contour depicted with a dashed white line) stained with anti-GFP (green). Scale bars, 50 µm. (**D**) RNAi-mediated downregulation of *Lgr3* (*Lgr3-IR*) in PIL/GCL neurons using *PIL/GCL>* rescues the delay induced by *Tub-dilp8(2)*. Box plot showing pupariation time (Time after egg laying (AEL)) in h of (N) larvae. As a negative control, a stock carrying *PIL/GCL>* was crossed with *w[1118]* animals. Whiskers extend to minimum to maximum value, each dot represents one individual, the vertical bars represent the median. *P* < 0.001, One-way ANOVA test. Genotypes sharing the same letter are not statistically different at α = 0.05, Tukey’s post-hoc test. **(E)** Downregulation of the *alpha subunit of the heterotrimeric Gs*, *Gɑplhas* (*Galphas-IR*; *Gɑs-IR*) in PIL/CGL neurons using the broad driver *R19B09>* rescues the delay induced by *Tub-dilp8(2)*. As negative controls (blue), stocks carrying either *R19B09>* or *Galphas>IR* were crossed with *w[1118]* animals. *P* < 0.001, One-way ANOVA test. Rest, as in (D). **(F)** Downregulation of the *Gɑplhas* (*Gɑs-IR*) in PIL/CGL neurons using the sparse driver *R19B09.3D>* rescues the delay induced by *Tub-dilp8(2)*. Box plot showing pupariation time (AEL) in h of (N) larvae. *P* < 0.001, One-way ANOVA test. Rest, as in (D).

To verify if the intersection strategy worked as expected, we tested if *PIL/GCL>* indeed drove expression in PIL/GCL neurons and not in VNC neurons. For this, we crossed *PIL/GCL>* animals with a stock carrying *UAS-myr::tdTomato; sfGFP::Lgr3*, and then performed immunohistochemistry of the F1 larval CNS to analyze the colocalization between sfGFP::Lgr3 and PIL/GCL>myr::tdTomato, detected with antibodies against GFP and myr::tdTomato (anti-DsRed), respectively. We expected to observe a colocalization of both reporters in the PIL/GCL neurons and limited or no expression in other cells across the CNS, especially in the *tsh* domain of the VNC. Consistent with our expectations, our results show clear colocalization between *sfGFP::Lgr3* and PIL/GCL>myr::tdTomato in 2.36 ± 0.77 PIL/GCL neurons per brain (*N* = 14), and limited and variable expression in other cells (**Figure 4B**). Importantly, in 11/14 brains analyzed, there was at least 1 PIL/GCL neuron labeled in both brain hemispheres. 1/14 had 2 ipsilateral PIL/GCL neurons labeled, and 2/14 had only 1 PIL/GCL neuron labeled. Based on these results alone, the PIL/GCL> intersectional driver is expected to behave similarly or better than the *R19B09.2>* driver, which despite only driving expression in a subset of PIL/GCL neurons (**Figures 2C-D**), significantly disrupts PIL/GCL neuron function upon RNAi or thermogenetic manipulations (**Figures 1D and 3C**). To test if the incomplete intersection was a characteristic of the *PIL/GCL>* stock or rather a limitation of the sensitivity of the colocalization assay (as reported above for the enhancer bashing experiments), we used the more sensitive *Lgr3-IR, sfGFP::Lgr3* assay, and expressed *Lgr3-IR* under the control of *PIL/GCL>* (*PIL/GCL>Lgr3-IR*) in the presence of *sfGFP::Lgr3*, followed by immunofluorescence assays, as described above. We found no detectable somatic sfGFP::Lgr3 expression in the neuroanatomical region corresponding to PIL/GCL neurons (**Figure 4C**). These results suggested that *PIL/GCL>Lgr3-IR* is sufficiently strong to reduce Lgr3 protein levels in PIL/GCL neurons to undetectable by immunofluorescence assays, similarly to the *R19B09_CRM>* drivers. Importantly, in other neuroanatomical regions, sfGFP::Lgr3 expression was not detectably affected (**Figure 4C**), confirming the specificity of the intersection for the PIL/GCL neurons.

Next, we used *PIL/GCL>* to functionally test the hypothesis that PIL/GCL neurons are the key neurons where *Lgr3* is required to transduce the Dilp8 developmental-delay signal. For this, we expressed *Lgr3* RNAi in *PIL/GCL>* neurons (*PIL/GCL>Lgr3-IR*) and scored the timing of pupariation in the presence of ectopic Dilp8 activity via the *Tub-dip8(2)* cassette and compared this to a series of controls (**Figure 4D**). As expected, *Tub-dilp8(2), PIL/GCL>Lgr3-IR* animals pupariated significantly before *Tub-dilp8(2)* animals or *Tub-dilp8(2), PIL/GCL>* without the *Lgr3-IR* cassette (*P <* 0.05, Tukey’s *post-hoc* test). The suppression of the delay was not as extensive as the effect of *MZ699>* alone (control *PIL/GCL>* without the *FRT-GAL80* cassette), but was statistically significant (*P <* 0.05, Tukey’s *post-hoc* test). This could be due to the stochastic nature of the recombinase-mediated genetic manipulation in larvae, so that not all PIL/GCL neurons would start expressing enough *Lgr3-IR* in time for a full biological effect. Nevertheless, taken together with the enhancer bashing results, these intersectional genetics experiments unequivocally demonstrate that *Lgr3* is required in the PIL/GCL neurons for the Dilp8-dependent developmental delay, thereby confirming this part of the model (**Figure 1A**).

### Lgr3 signaling in PIL/GCL neurons

#### Stimulatory G protein is required in PIL/GCL neurons for proper pupariation

Dilp8-dependent activation of Lgr3 in PIL/GCL interneurons is associated with an increase in the intracellular levels of the second messenger cyclic adenosine monophosphate (cAMP) [11,20,21] (**Figure 1C**). However, the requirement of cAMP-dependent signaling for the Dilp8 and Lgr3-dependent activities has never been formally tested. G-protein-coupled receptors increase intracellular cAMP levels by activating the heterotrimeric stimulatory G protein G_s_, which then activates adenylyl cyclase to increase intracellular cAMP levels and stimulate the cAMP-dependent signaling pathway [45]. To demonstrate that Dilp8-dependent Lgr3 signaling in PIL/GCL neurons requires heterotrimeric G_s_, we first used RNAi to knock down the *alpha subunit of the heterotrimeric G_s_, Gɑ_s_ (Gɑ_s_-IR*) in PIL/CGL neurons using the *R19B09>* driver (*R19B09>Gɑ_s_-IR*) in animals carrying the developmental delay-causing *Tub-dilp8(2)* transgene, and quantified the timing of pupariation of these animals and of controls. We found that *R19B09>Gɑ_s_-IR* completely suppressed the *Tub-dilp8(2)-*dependent delay in the onset of metamorphosis (**Figure 4E**). Unfortunately, our efforts to knock-down *Lgr3* with greater specificity in PIL/GCL neurons using the intersectional genetics strategies described above were not fruitful due to synthetic lethality of the system with the *Gɑ_s_-IR* construct insertion. To overcome this limitation, we resorted to *R19B09.3D>*, one of the sparse PIL/GCL*-*neuron drivers identified in the enhancer bashing experiments (**Figure 1D and 2E**). Accordingly, *R19B09.3D>Gɑ_s_-IR* strongly suppressed the *Tub-dilp8(2)*-dependent developmental delay (**Figure 4F**). These results are consistent with previous findings [20,21] and strongly support a requirement for a *Gɑ_s_* and hence a cAMP-dependent transduction pathway in PIL/GCL neurons for imaginal disc growth coordination during early larval development.

### Neurogenetic modulation of PIL/GCL activity

#### PIL/GCL neuron silencing suppresses the Dilp8-dependent developmental delay

We next set out to test another critical assumption of the Dilp8-Lgr3 developmental stability pathway model: that Lgr3 activation by Dilp8 leads to PIL/GCL neuron activation (**Figure 1A**). Typically, neuronal activation consists of membrane depolarization followed by synaptic vesicle release at axon terminals [46]. To test if these processes are required in PIL/GCL neurons downstream of Dilp8-Lgr3 signaling, we used a neurogenetic strategy. We first hyperpolarized PIL/GCL neurons using the *PIL/GCL>* line coupled with a *UAS-FRT-stop-FRT-Kir2.1::GFP* construct (which expresses GAL4-dependent *Kir2.1::GFP* upon FLP-mediated removal of the stop cassette [47]) in the presence or not of *Tub-dilp8(3)* (a 3rd chromosome insertion of *Tub-dilp8* [21]) and scored for the pupariation timing. The hypothesis was that Kir2.1-induced hyperpolarization would counteract the effects of Dilp8/Lgr3-triggered PIL/GCL-neuron depolarization, thereby suppressing the delay caused by the presence of ectopic Tub-dilp8(3) expression. In agreement with our hypothesis, we found that *Tub-dilp8(3), PIL/GCL>Kir2.1::GFP* animals pupariated significantly earlier than *Tub-dilp8(3); UAS-FRT-stop-FRT-Kir2.1::GFP* controls without *PIL/GCL>* (*P <* 0.05, Tukey’s *post-hoc* test) (**Figure 5A**), demonstrating that *PIL/GCL>* neuron hyperpolarization suppresses the effect that Dilp8 has on PIL/GCL neurons. These results indicate that Dilp8-dependent Lgr3 signaling in PIL/GCL neurons leads to their depolarization and that this depolarization is required for the pupariation delay.

**Figure 5.**
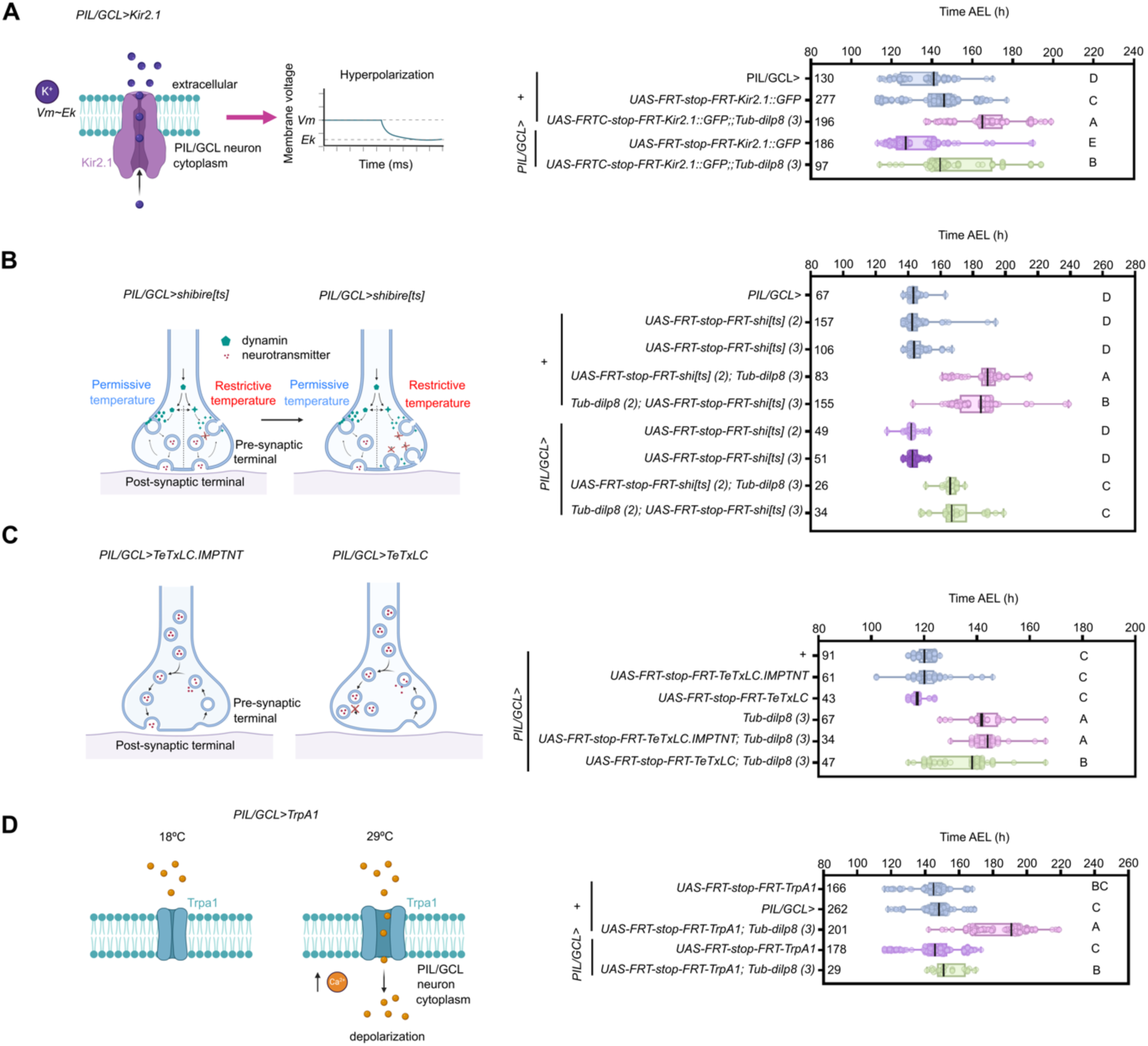
PIL/GCL neuron silencing or activation suppresses the Dilp8-dependent developmental delay. **(A)** Kir2.1-induced hyperpolarization in PIL/GCL neurons partially suppresses the developmental delay caused by the presence of ectopic Tub-dilp8 expression. Box plot showing pupariation time (Time after egg laying (AEL)) in h of (N) larvae expressing *Tub-dilp8(3)* and *PIL/GCL>Kir2.1* (green). Negative controls (blue), *PIL/GCL>* or *UAS-FRT-stop-FRT-Kir2.1::GFP* crossed with *w[1118]*. **(B)** Inhibiting synaptic transmission by blocking vesicle endocytosis using the temperature-sensitive dynamin mutant allele *shi[ts]* in PIL/GCL neurons partially suppresses the delay caused by the presence of ectopic *Tub-dilp8* expression. Box plot showing pupariation time (AEL) in h of (N) larvae expressing *Tub-dilp8(2) or Tub-dilp8 (3)* and *shi[ts]* (via constructs *UAS-FRT-stop-FRT-shi[ts](2)* or *UAS-FRT-stop-FRT-shi[ts](3)*) under control of *PIL/GCL>* (green). Negative controls (blue), *PIL/GCL>* or *shit[ts]* constructs crossed with *w[1118]*. **(C)** Inhibiting synaptic transmission by blocking vesicle exocytosis using the tetanus toxin light chain (TeTxLC) in PIL/GCL neurons partially suppresses the delay caused by the presence of ectopic Tub-dilp8 expression. Box plot showing pupariation time (AEL) in h of (N) larvae expressing *Tub-dilp8(3)* and *PIL/GCL>TeTxLC* (green). Negative controls (blue), *PIL/GCL>* crossed with *w[1118]*. Positive controls (pink), *PIL/GCL>* crossed with *Tub-dilp8(3)* or with animals carrying a mutated, inactive form or *TeTxLC, TeTxLC.IMPTNT*. **(D)** Thermogenetic activation of PIL/GCL neurons suppresses the Tub-dilp8 delay. Box plot showing pupariation time (AEL) in h of (N) larvae expressing *Tub-dilp8(3)* and *PIL/GCL>TrpA1* (green). Negative controls (blue), *PIL/GCL>* or *UAS-FRT-stop-FRT-TrpA1* crossed with *w[1118]*. In all panels, whiskers extend to minimum to maximum value, each dot represents one individual, the vertical bars represent the median. *P* < 0.0001, One-way ANOVA test. Genotypes sharing the same letter are not statistically different at α = 0.05, Tukey’s post-hoc test.

If PIL/GCL neurons depolarize upon Lgr3 activation, as the results above suggest, then it is a fair assumption–based on typical neuronal properties–that this depolarization is followed by the release of neurotransmitter-containing vesicles at the presynaptic terminals of PIL/GCL neurons. To test this, we blocked synaptic transmission in PIL/GCL neurons using two independent neurogenetic strategies: i) synaptic vesicle exhaustion by blocking vesicle endocytosis using the temperature-sensitive dynamin mutant allele, *shibire[ts] (shi[ts])* [48], and ii) direct synaptic vesicle exocytosis blocking using the tetanus toxin light chain (TeTxLC), which inhibits vesicle-membrane fusion events by cleaving members of the synaptobrevin/vesicle-associated membrane protein family [49]. In the first strategy, animals expressing *shi[ts]* in PIL/GCL neurons using two independent constructs (*PIL/GCL>UAS-FRT-stop-FRT-shi[ts](2)* or -*shi[ts](3)* [50]) in the presence of *Tub-dilp8(3)* or *Tub-dilp8(2)*, respectively, pupariated significantly earlier than their respective control *PIL/GCL>, Tub-dilp8(2 or 3)* animals (*P <* 0.05, Tukey’s *post-hoc* test) (**Figure 5B**), but also significantly later than negative controls not carrying the *Tub-dilp8(2 or 3)* constructs (*P <* 0.05, Tukey’s *post-hoc* test) (**Figure 5B**). This indicated that blocking synaptic transmission in PIL/GCL neurons partially suppresses the effect that Dilp8 has on PIL/GCL neurons, similarly to the results of neuronal hyperpolarization. This finding is highly consistent with the model that synaptic transmission occurs downstream of Dilp8-Lgr3 signaling in PIL/GCL neurons. In the second strategy, *Tub-dilp8(3)* animals expressing TeTxLC in PIL/GCL neurons using a similar genetic approach as above (*Tub-dilp8(3), PIL/GCL>UAS-FRT-stop-FRT-TeTxLC*), also pupariated significantly earlier than the positive controls expressing a mutated, inactive form of TeTxLC, TeTxLC.IMPTNT (*Tub-dilp8(3), PIL/GCL>UAS-FRT-stop-FRT-TeTxLC.IMPTNT*) or not expressing any TeTxLC version (*Tub-dilp8(3), PIL/GCL>*) (*P <* 0.05, Tukey’s *post-hoc* test for both conditions) (**Figure 5C**), but again later than the negative controls not carrying the *Tub-dilp8(3)* cassette (*P <* 0.05, Tukey’s *post-hoc* test for all conditions) (**Figure 5C**). These experiments demonstrate that expression of TeTxLC in PIL/GCL neurons partially rescues the Dilp8-dependent delay in pupariation timing, and are highly consistent with the *Kir2.1* and *shit[ts]* experiments described above. As the three neuronal silencing strategies rely on the same driver intersection, and in the three cases the rescue of the *Tub-dilp8(2 or 3)*-effect was partial, rather than complete, the most conservative conclusion of these experiments is that PIL/GCL-neuron depolarization and synaptic transmission are required for the Dilp8/Lgr3-dependent delay, confirming another important assumption of the model.

Interestingly, in the absence of ectopic Tub-Dilp8 expression, all neuronal silencing experiments led to small anticipations of the median time of pupariation, although this only reached statistical significance for the neuronal hyperpolarization experiments. Namely, *PIL/GCL>Kir2.1::GFP* animals pupariated slightly earlier than *PIL/GCL>* or *UAS-FRT-stop-FRT-Kir2.1::GFP* negative controls (*P <* 0.05, Tukey’s *post-hoc* test) (**Figure 5A**). In contrast, the difference was not statistically significant when *shi[ts]* or *TeTxLC* were expressed in PIL/GCL neurons in the absence of *Tub-dilp8* (*P >* 0.05, Tukey’s *post-hoc* test) (**Figure 5C**). Hence, it is clear that in the experimental conditions used in this study, this small effect is not robust to the different neurogenetic silencing methods. However, the absence of a statistically significant effect in the synaptic transmission manipulation experiments could be due to the relatively small effect size of this anticipation (a few hours). Larger sample sizes and/or other assay setups, such as those using larvae synchronized at the L2-L3 molt, might be more appropriate to detect such smaller effects. For instance, such assays were used to demonstrate that *Lgr3* mutation alone leads to a ∼4-h anticipation of the pupariation time in control conditions (*i.e.,* in the absence of any genetic or environmental challenge) [20]. Future studies can address this interesting question, because a minor anticipation of pupariation time due to PIL/GCL neuron silencing would also be consistent with the model where PIL/GCL neuron activity prolongs the larval phase, in this case supposedly in response to endogenous Dilp8-Lgr3 signaling.

#### PIL/GCL neuron activation by depolarization is not sufficient to delay the onset of pupariation

According to the current model (**Figure 1A and C**) and to the neuronal silencing results described above, the neurogenetic activation of PIL/GCL neurons should or could lead to a delay in the larval-to-pupal transition, even in the absence of any endogenous or exogenous stimuli that activates the Dilp8-Lgr3 pathway. However, our enhancer bashing experiments suggested that thermogenetic PIL/GCL-neuron activation using TrpA1 was not sufficient to cause a pupariation timing delay (see *R19B09.3A>TrpA1* in **Figure 3C-D**). To further support these findings, we repeated the *TrpA1* thermogenetic activation experiments, but now using the same intersectional genetics strategy used for the neuronal silencing experiments–namely, by coupling *PIL/GCL>* with a *UAS-FRT-stop-FRT-TrpA1* construct [51] (*PIL/GCL>UAS-FRT-stop-FRT-TrpA1*). Importantly, *PIL/GCL>UAS-FRT-stop-FRT-TrpA1* animals also did not pupariate significantly later than controls carrying either the *PIL/GCL>* or *UAS-FRT-stop-FRT-TrpA1* cassettes alone (*P* > 0.05, Tukey’s *post-hoc* test) (**Figure 5D**). Therefore, two independent techniques and drivers indicate that favoring PIL/GCL neuron depolarization via increased cation influx is not sufficient to mimic the behavioral effects of the Dilp8-dependent activation of Lgr3 in these cells. Hence, whereas the neuronal silencing experiments demonstrate that PIL/GCL neuron depolarization is necessary for the Dilp8-dependent delay in the larval-to-pupal transition, we can now conclude that PIL/GCL neuron depolarization alone is clearly not sufficient for this delay to take place.

#### PIL/GCL neuron activation by depolarization suppresses the Dilp8-dependent developmental delay

We next wondered if this thermogenetically-induced depolarization of PIL/GCL neurons would increase the developmental delay effect in sensitized animals, which already carry a delay-inducing *Tub-dilp8(3)* cassette, and thus supposedly have activated Lgr3 signaling in PIL/GCL neurons. However, inconsistently with this expectation, results showed that thermogenetic activation of PIL/GCL neurons did not further extend the Dilp8-dependent delay (**Figure 5D**). Instead, to our surprise, it suppressed it (*P <* 0.05, Tukey’s *post-hoc* test against *UAS-FRT-stop-FRT-TrpA1*; *Tub-dilp8(3)* animals) back to the levels of *PIL/GCL>* controls (*P >* 0.05, Tukey’s *post-hoc* test) (**Figure 5D**). The simplest interpretation of these results is that neuronal silencing is also necessary for the Dilp8-dependent delay in pupariation. Hence, apart from rejecting the current PIL/GCL functional model (**Figure 1A and C**), as described above, these results further suggested that Dilp8 activation of Lgr3 in PIL/GCL neurons is partially inhibitory, instead of only excitatory, and that this inhibition can be counteracted by cation influx to the PIL/GCL to significantly anticipate pupariation timing [20].

An alternative explanation for the TrpA1 effect would be that chronic TrpA1 activation negatively interferes with some critical function of PIL/GCL neurons, reducing their activity instead of activating them. While it is difficult to exclude this possibility completely, we believe this is unlikely the case for PIL/GCL neurons for three main reasons. First, temperature-dependent TrpA1 activation in larval neurons (for instance, motor neurons or serotonergic neurons) has been shown to increase neuronal activity as expected and in the case of motor neurons drive tonic action potentials with little adaptation [37-41]. Second, whereas TrpA1 is constitutively expressed in the PIL/GCL neurons, in our experiments it is only activated by the temperature shift at the end of L2/beginning of L3 stage, so the time period is relatively short - even more so if one considers that the sensitive time for Dilp8 activity and hence PIL/GCL neuron activity is thought to occur most prominently during the first half of the 3rd instar larval phase before the mid-third instar transition [11,13,20]. Third, circuit architecture is mostly stable throughout the larval stages [52], so that primary neurons, such as PIL/GCL neurons, are expected to establish their connections by the L1 stage, maintaining their overall shape and connectivity up to pupariation. To confirm this is the case for PIL/GCL neurons, we studied their neuroanatomy by direct immunofluorescence observation during development using the sparse *R19B09.3D>* driver, and found no obvious difference in gross PIL/GCL neuron morphology from L1 to Late L3 (**Supplementary Figure S3**). Hence, TrpA1 activation at late L2/early L3 is not expected to interfere with PIL/GCL neuron growth or synaptogenesis.

How can the same pair of bilateral neurons require both silent and active phases to exert their biological effects? These apparently opposing results can be reconciled in a model where the activation of Lgr3 by Dilp8 in PIL/GCL neurons leads to oscillations in membrane potential, such that PIL/GCL neurons start transitioning between depolarized and hyperpolarized states. Such phenomena can arise by two main mechanisms, via neurons with intrinsic oscillating properties, the so-called pacemaker or pacemaker-like neurons, or via simple feedback inhibition circuits from neurons which do not have intrinsic bursting properties (reviewed in [29-30]). In the case of PIL/GCL neurons, the neurons would acquire such activity following Lgr3 activation by Dilp8. In a model where PIL/GCL neurons acquire intrinsic pacemaker activity upon Lgr3 activation, this would be analogous to how neurons in the rodent developing superficial dorsal horn neurons and adult deep dorsal horn neurons, for instance, switch between pacemaker and non-pacemaker activity upon stimulation by different factors [53-54], in the latter case due to G-protein coupled receptor-dependent activation of inward-rectifying potassium (Kir) conductance [53]. Similarly, oxytocin/vasopressin-related neuropeptides have long been known to induce bursting pacemaker currents in mollusk neurosecretory and central neurons [55,56]. Alternatively, in a model where PIL/GCL neurons do not acquire intrinsic pacemaker activity, they could acquire oscillatory behavior via the Lgr3-dependent network property changes, such as by synaptogenesis, for instance, or any other mechanism that activates reciprocally inhibitory synapses between themselves and confer the circuit the potential for feedback inhibition [30].

### Intrinsic and network properties of PIL/GCL neurons

#### PIL/GCL neurons express Cyclin A, a cell cycle protein involved in sleep time regulation in postmitotic neurons

To start gaining insight into PIL/GCL neuron responses, we first tested whether PIL/GCL neurons were involved in the circadian pacemaker pathway, which controls many self-sustained rhythms in physiology and behavior with ∼24 h periodicity [57]. L3 larvae contain 6-8 clock neurons, which are positive for the gene *period* (*per*) [58], a product of the transcription factors Clock and Cycle (reviewed in [59]). Even though the localization of PIL/GCL neurons is not totally consistent with that of known clock neurons, we tested whether *Lgr3* would be required in *per*-positive cells, by knocking down *Lgr3* in *per-GAL4* cells [60] (*per>Lgr3-IR*) and asking if this would suppress the *Tub-dilp8-*dependent delay in development. If yes, this would mean that PIL/GCL neurons are clock neurons. Our results were clearly negative, showing that *Lgr3* is not required in *per>-*positive cells, which indicates that PIL/GCL neurons are not clock neurons (**Figure 6A**). The neuropeptide pigment-dispersing factor (Pdf) is expressed in a subset of clock neurons [61], and is a major regulator of neuronal clock output, by modulating Pdf receptor (Pdfr)-positive cells [62]. If PIL/GCL neurons could be entrained by Pdf, they would be Pdfr-positive. Hence, we tested this by knocking down *Lgr3* in *Pdfr(B)-GAL4* cells [63] (*Pdfr>Lgr3-IR*) and asking if this was sufficient to rescue the *Tub-dilp8* delay. Again, our results were negative (**Figure 6A)**, showing that *Lgr3* is not necessary in *Pdfr>-*positive cells for the developmental stability pathway, and indicating that PIL/GCL neurons are not likely not directly Pdf-responsive, or directly entrained by the clock neurons. Hence, it is likely that PIL/GCL neurons utilize other mechanisms of rhythmicity or membrane potential fluctuation.

**Figure 6.**
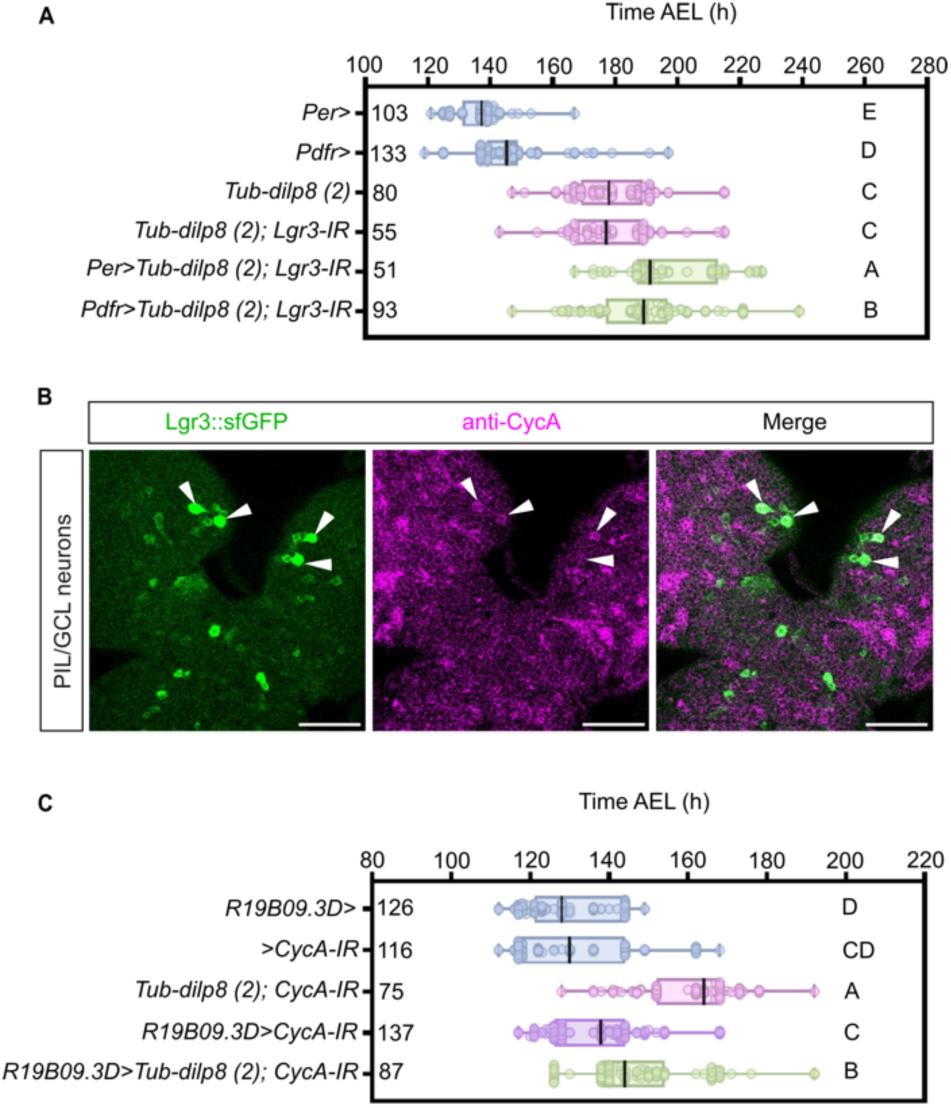
Lgr3 is not required in circadian pacemaker cells, and PIL/GCL neurons express and require CycA. **(A)** Downregulation of the *Lgr3* (*Lgr3-IR*) in *Per>* or *Pdfr>* cells does not rescue the delay induced by *Tub-dilp8(2)*. Box plot showing pupariation time (Time after egg laying (AEL)) in h of (N) larvae. As negative controls, stocks carrying either *Per>* or *Pdfr>* were crossed with *w[1118]* animals. Whiskers extend to minimum to maximum value, each dot represents one individual, the vertical bars represent the median. *P* < 0.001, One-way ANOVA test. Genotypes sharing the same letter are not statistically different at α = 0.05, Tukey’s post-hoc test. **(B)** Depicted is the sum of confocal z-stack slices of an immunohistochemistry preparation obtained from a dissected 3rd instar larva CNS stained with anti-GFP (sfGFP::Lgr3, green) and anti-CycA (magenta). The anterior/frontal CNS region is shown, with four PIL/GCL neuron stomata (arrowheads, strong green) highlighted, of which three are clearly positive for elevated levels of CycA (magenta).). Scale bars, 50 µm. **(C)** Downregulation of the *CycA* (*CycA-IR*) in *R19B09.3D>* cells partially rescues the delay induced by *Tub-dilp8(2)*. Negative controls (blue), *R19B09.3D>* or *>Cyc-IR* crossed with *w[1118]*. This experiment was conducted in parallel with Figures 4F and Supplementary Figure S4E, so that *R19B09.3D>* animals are the same in the three figures panels. *P* < 0.001, One-way ANOVA test. Rest, same as (A).

In cycling cells, cyclins form a complex with a catalytic subunit, cyclin-dependent kinase (CDK), to participate in cell-cycle-linked oscillations [64-66]. Cyclin A (CycA), which is widely known for its roles in mitosis [64-67], is also expressed in postmitotic neurons in adult flies, where it functions in a dedicated, yet still poorly-understood pathway to determine sleep length [27]. Adult CycA-positive neurons are frequently, but not always, intermingled with circadian pacemaker neurons in the adult brain. However, the sleep promoting function of CycA, and of other players in the same pathway [namely, Regulator of Cyclin A1 (Rca1) and TARANIS (TARA), which are *Drosophila* homologs of the mammalian early mitotic inhibitor (Emi1/F-box only protein 5, FBXO5) and the Trip-Br (SERTAD) family of transcriptional coregulators, respectively] is independent of the circadian system [27,28]. To test whether PIL/GCL neurons are involved in a similar pathway, we stained larval CNS preparations from *sfGFP::Lgr3* animals with an antibody against full length *Drosophila* CycA. As expected, CycA stains several cycling cells in the larval brain, including neuroblasts, their lineages, and inner and outer proliferating centers in the optic lobes (**Supplementary Figure S4**). Surprisingly, PIL/GCL neuron stomata stained positive for CycA (**Figure 6B**). Another pair of Lgr3-positive neurons, the Midline Internal Lgr3-positive (MIL) neurons, required for proper progression of the pupariation motor program [68], also stained positive for CycA (**Supplementary Figure S4**). These results suggest that PIL/GCL neurons and maybe other Lgr3-positive neurons might use similar time-setting mechanisms as sleep pathway neurons.

To genetically define a requirement for CycA in PIL/GCL neurons, we knocked-down *CycA* (*CycA-IR*) in PIL/GCL neurons using *R19B09.3D>* in the presence of *Tub-dilp8(2)*. *R19B09.3D>CycA-IR* animals pupariated significantly earlier than *Tub-dilp8(2)* animals, but also later than controls (**Figure 6C**), showing that *CycA* RNAi partially suppresses the *Tub-dilp8* delay to a similar extent as PIL/GCL neuron silencing or activation (*i.e.,* oscillation) (**Figure 5**). Interestingly, knockdown of the CycA-regulator *Rca1* in *R19B09.3D>* cells had no effect on the *Tub-dilp8(2)* delay (**Supplementary Figure S4**), suggesting that the role of CycA in PIL/GCL neurons is not exactly the same as during sleep length regulation, where Rca1 is also required [27,28]. The role of CycA in larval growth-coordinating neurons adds to an increasing list of postmitotic roles for cyclins in animals [69-77]. Apart from sleep-length, cyclins can regulate synaptic plasticity via modulation of the activity of postmitotic-neuron CDKs, such as CDK5-like proteins, in vertebrates and invertebrates, and regulate centriole amplification in multiciliated cells in vertebrates. Alternatively, CycA could also be involved in endoreduplication cycles (or “endocycles”; G/S phase oscillations that skip mitosis), even though there is no evidence that PIL/GCL neurons–or any other larval neuron at this stage–undergo endocycles, since the brains of newly eclosed adult flies are 98–99% diploid [78]. Whichever the mechanism, the requirement of CycA for proper PIL/GCL function, and its expression in other Lgr3-positive neurons, connects cyclins, which participate in well-described oscillating mechanisms in cycling cells, to putatively oscillating larval neurons and indicate an intimate relationship between the relaxin pathway and cyclin expression in postmitotic neurons.

#### PIL/GCL neurons consist of two distinct neurons, a local (PIL/GCL-LN) and a projection (PIL/GCL-PN) neuron

Rhythmically oscillating neuronal networks can arise from either intrinsic properties or network properties (reviewed in [29,30]). To start gaining insight into possible network properties of the PIL/GCL neurons, we first characterized PIL/GCL neuron neuroanatomy in more detail using MultiColor FlpOut (MCFO) [79] to stochastically label *R19B09.3D>-*positive neurons, which include the PIL/GCL neurons (**Figure 7A-C**). Briefly, the MCFO strategy used (*R19B09.3D>MCFO*) included, in addition to *R19B09.3D*>, a panneuronally-driven FLPL (*R57C10-FLPL*), and a series of *UAS-* and FLPL-dependent reporters (*10xUAS(FRT.stop)myr::smGFP-HA, 10xUAS(FRT.stop)myr::smGFP-V5, 10xUAS(FRT.stop)myr::smGFP-FLAG*), which we detected using an anti-V5 antibody and an anti-sfGFP antibody [which labels smGFP (“spaguetti-monster GFP”) in all *R19B09.3D>*-positive FlpOut clones]. We consistently found clones with two different branching patterns originating from cell bodies placed in the PIL/GCL neuron region (anterior/rostral-medial region of the CNS). Namely, while both PIL/GCL neurons send bilateral (both ipsi- and contralateral) anterior projections (**Figures 7A**, arrows) to the same neuropil compartments–crossing the midline only once–, only one PIL/GCL neuron from each hemibrain has an additional ipsilateral descending projection to the SEZ (**Figures 7B-C**, arrows on descending projection). This latter PIL/GCL neuron can be classified both as a projection neuron (PN, which are interneurons that carry information between brain regions; following the classification of [80], which does not distinguish between first-hop PNs and output neurons) and also a descending neuron towards the SEZ (DN-SEZ). We named this neuron PIL/GCL-PN. The other PIL/GCL neuron has a branching pattern consistent with a local neuron (LN), defined as a neuron that processes information within a single neuropil compartment (ipsilaterally, contralaterally, or bilaterally). Hence, we called this neuron, PIL/GCL-LN. PIL/GCL-LNs are most likely of the bilateral type (**Figure 7A**), which would make it one of the eight known bilateral LNs in the larval brain [80]. This should facilitate their identification in the reconstituted L1 electron microscopy data for further connectome studies.

**Figure 7.**
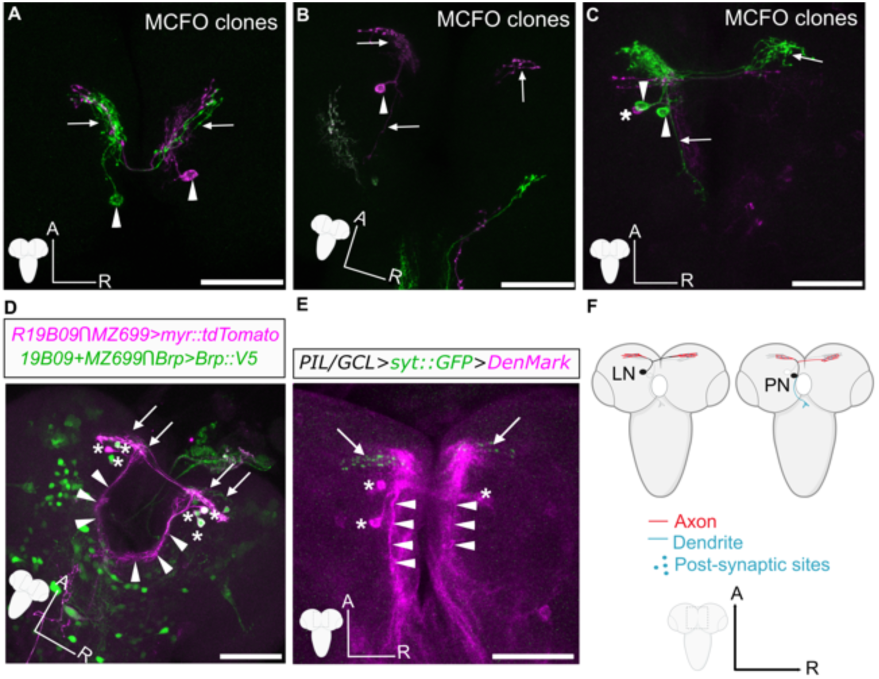
PIL/GCL neuroanatomy. **(A-C)** Sum of confocal z-stack slices of MCFO labeling of L3 larval stage neurons with panneuronally (*R57C10>*)-driven expression of *FLP* and *R19B09.3D>-*driven expression *UAS-*driven MCFO markers differentially labeled here with anti-GFP (green) and anti-V5 (magenta). Arrowheads, PIL/GCL neuron cell bodies. Arrows, PIL/GCL neurons projections. Asterisk, a #5 neuron soma. **(A)** Two MCFO clones showing two apparently homologous PIL/GCL local neurons (PIL/GCL-LNs, magenta and green). **(B)** A single PIL/GCL projection neuron (PIL/GCL-PN) clone (magenta). This neuron has similar ipsilateral and contralateral projections to the PIL/GCL-LNs depicted in (A), but in addition, has a descending projection towards the SEZ. **(C)** Two ipsilaterally-localized PIL/GCL neuron clones are labeled in green, but only a single green-labeled projection descends towards the SEZ. Hence, these two PIL/GCL neurons represent a PIL/GCL-LN and a PIL/GCL-PN. Another neighboring #5 neuron (magenta), which has a different pattern of ipsi- and contralateral projections (more posterior and straight) than those of PIL/GCL neurons is also visible in this image. It also has a descending projection towards the SEZ. **(D)** Presynaptic sites of #5 neurons, a group of neurons which include the PIL/GCL neurons, are almost exclusively located in the anterior bilateral projections. Shown is the sum of confocal z-stack slices of dissected wandering L3 larval stage CNSs stained with anti-GFP (green). In this preparation, *R19B09 ∩ MZ699>* drives expression of *myr::tdTomato* (magenta) and *R19B09>>* and *MZ699>* also independently unlock the expression of *brp*-driven (neuronally-driven) Brp::V5 (detected with an anti-V5 antibody, green) in R19B09+Brp+ or MZ699+Brp+ neurons, respectively. This is possible as in this cross there is an *8xLexAop2-FLPL* and a *UAS-FLP* cassette, either of which can mediate FRT removal of the *brp-FRT-stop-FRT-V5* cassette. Hence, all *MZ699>-*positive #5 neurons are labeled. Asterisks, #5 neuron somata. Arrows, anteriormost PIL/GCL neuron projections (green). Arrowheads, descending projections. **(E)** Dendritic and axonal compartments of PIL/GCL neurons labeled with somatodendritic marker, DenMark (magenta), and the synaptic vesicle marker, syt::GFP (green), respectively. PIL/GCL neuron (and possibly other #5 neuron) cell bodies (asterisks), axonal compartments (arrows), and dendritic compartments (arrowheads) are depicted. Notice the absence of syt::GFP in the descending projections, indicating they are mostly, if not exclusively enriched for postsynaptic sites (dendritic). A full CNS confocal projection of this preparation is depicted in **Supplementary Figure S6**. **(F)** Model describing PIL/GCL neuron neuroanatomy. Circles represent the cell body of each PIL/GCL neuron (PIL/GCL-LN and -PN, for local and projection neuron, respectively), which send bilateral anterior projections to the homologous neuropil compartments. Only PIL/GCL-PN also sends a descending projection to the subesophageal zone (SEZ), localized below the esophageal foramen. Projections are colored to depict the axonal (red) and dendritic (blue) compartments of both cell types. We hypothesize that the anterior projections of both PIL/GCL-LNs and -PNs are bilateral in nature, with bilateral presynaptic and postsynaptic sites, but this remains to be formally tested with connectome data. Scale bars, 50 µm.

Other brain neurons were also labeled by *R19B09.3D>MCFO*, albeit less frequently, including one or more neurons that we believe could be other #5 neurons, with cell bodies located in the same anatomical region as the PIL/GCL neurons cell bodies (**Figure 7C, asterisk**), and another more posterior neuron (**Supplementary Figure S5**). The other #5 neuronal type projects to both sides and to the SEZ, similarly to the PIL/GCL-PN, but has a more straight/linear path in the mediolateral axis. The PIL/GCL neurons can be distinguished by their more curvilinear projections to a more anterior area of this neuropil compartment (**Figure 7C, green**). Whereas the full relevance of these neuroanatomical findings are not completely clear at this stage, we can already learn at least two important facts that are relevant to this study: first, the different branching patterns between PIL/GCL-LN and -PNs reveals an unforeseen layer of complexity in PIL/GCL neuron biology, suggesting specialized functions for each PIL/GCL neuron subtype, rather than a single function as previously modeled. Second, both PIL/GCL neurons (LN and PN) seem to share reciprocal bilateral projections, suggesting that they might be subject to contralateral feedback, which is a common oscillator-triggering mechanism [30].

#### PIL/GCL-LN and -PN neurons are bilateral and likely form reciprocal loops

To gain more insight into the directionality of information processing of PIL/GCL neurons, we first expressed a V5-tagged version of the pre-synaptic active-zone marker, Bruchpilot [81] (Brp::V5), in *R19B09* ∩ *MZ699>* neurons using a similar FlpOut strategy [82]. Because of the particularities of this cross, we see Brp::V5 from both R19B09 *and* MZ699 drivers, which includes all #5 neurons. Nevertheless, we found strong Brp-labeling in the anterior projections of the PIL/GCL neurons (and other #5 neurons), but no or very limited positive staining in their descending projections (**Figure 7D**). The simplest interpretation of this finding is that the descending branches of all #5 PNs, which include the PIL/GCL-PNs, are mostly, if not exclusively dendritic, as they are largely devoid of Brp staining. To independently confirm this finding, we performed specific co-labeling of PIL/GCL neurons with pre- and postsynaptic markers. Namely, we used the synaptic vesicle marker, synaptotagmin 1 (syt::GFP) (*UAS-syt.eGFP*; ref.[83]), and the somatodendritic marker, DenMark (<u>Den</u>dritic <u>Mark</u>er, a hybrid protein of the mouse dendritically-targeted protein, ICAM5/Telencephalin, and mCherry; ref.[84]), which are expected to label most clearly the PIL/GCL neuron axonal and dendritic compartments, respectively. In agreement with the experiments described above, we found that the PIL/GCL neuron anterior projections (and likely those of other neighboring #5 neurons) are decorated strongly with syt::GFP, whereas DenMark staining was found most prominently, but not exclusively, in their soma and descending projections (**Figure 7E**). These results confirm that the anterior bilateral projections of PIL/GCL neurons (both LNs and PNs) contain their axons (and relatively less postsynaptic markers), and that the descending ipsilateral projection of the PIL/GCL-PN is mostly or exclusively dendritic. We conclude that both PIL/GCL neuron types, -LNs and -PNs, send anterior bilateral projections towards homologous compartments. The LNs (and DNs) likely transmit and receive information in this neuropil region, likely forming reciprocal contralateral loops, which are enriched in brain areas that play a role in action-selection and learning [80]. On the other hand, only the PIL/GCL-PN dendrites collect ipsilateral information from more posterior brain regions and the SEZ– a zone receiving rich sensory information input [80]. Connectome data from the electron microscopy-based reconstruction of the CNS of an L1 larva show that there are at least 566 PNs with a similar neuroanatomy in the cluster of brain neurons with ipsilateral dendrites and bilateral axons [80]. Only a subset of these have cell body localizations and branching patterns consistent with PIL/GCL-PNs, which should facilitate their identification in the L1 connectome dataset. A model summarizing the neuroanatomical results for each PIL/GCL neuron is described in **Figure 7F**.

### A REVISED MODEL OF PIL/GCL NEURON ACTIVITY IN THE DEVELOPMENTAL STABILITY PATHWAY

#### PIL/GCL neurons as a model for hypothalamic-like multimodal integration

While further work is required to match PIL/GCL neuron FlpOut-clone neuroanatomy to the reconstructed electron microscopy synaptic-level pattern, which is critical to describe the full list of its upstream and downstream circuit components, our findings nevertheless open up the possibility that one of the functions of PIL/GCL neurons is to integrate SEZ-derived primary sensory inputs with the endocrine tissue-damage Dilp8 signal, to temporarily block the progression of pupariation and promote developmental stability. PIL/GCL neurons are therefore in a privileged position to integrate external and internal state information, which is the basis of encoding adaptive behavior [85]. Interestingly, such integration, which is similar in concept to neuronal binding (*i.e.,* how items encoded in distinct brain circuits or neural populations can be combined for perception, decisions, and actions [86]), is one of the functions proposed for oscillatory neurons or circuits [86-89]. PIL/GCL neurons are also localized in a fly brain region orthologous to the mammalian hypothalamus [90], which performs similar multimodal integrations of sensory, metabolic, visceral, and hormonal signals, being a key brain center modulating transitions in innate behaviors and states, many times via changes in oscillatory behavior of hypothalamic neurons [29,91-94]. For example, certain hypothalamic neurons possess a hyperpolarization-activated current, the h-current (also termed the ‘pacemaker current’), which also produces the heart beat [95]. It will be interesting to verify if PIL/GCL neurons also possess such properties. Whereas we present neurogenetic and neuroanatomical evidence consistent with oscillatory behavior, one limitation of our study is that in the absence of neurophysiological observations, it is unclear what type of oscillatory behavior these neurons would acquire upon Dilp8 stimulation and for how long they maintain it. As discussed above, our data could be explained either by intrinsic or network properties. But also, within the realm of intrinsic properties, pacemaker-like neurons can have wildly different intrinsic membrane properties, such as endogenous bursting and plateau potentials, which originate in bistable neurons that enter and leave a plateau when receiving a depolarization and hyperpolarization pulse, respectively [96]. Our findings for PIL/GCL would be consistent with either type of intrinsic membrane property.

#### PIL/GCL neurons as a model for neuropeptide-activated oscillatory behavior in higher brain areas

As mentioned above, the neuroanatomy of PIL/GCL neurons is consistent with that of other spontaneously oscillating circuits. This is most notable for certain central pattern generators, whose rhythmicities are well-known to be activated, modified, and terminated by neuromodulators [30], but also of other rhythmically-active neuronal networks generating non-rhythmic behaviors, such as sleep, arousal, and other higher brain functions (reviewed in [29]). The oscillating hypothalamic neurons described above, for instance, are affected by the sleep-modulating peptide, cortistatin [95]. Therefore, our key finding that Dilp8 triggers both depolarized and hyperpolarized states in PIL/GCL neurons is likely of broad relevance and can serve as a model for studying these responses in higher brain circuits.

It is also relevant that our work points to an involvement of the Dilp8-Lgr3 relaxin-like pathway in the control of rhythmically active neuronal networks. Interestingly, in vertebrates, a Dilp8 orthologue, relaxin 3, and its receptor [relaxin-family peptide receptor 3 (RXFP3)] in the medial septum complex are involved in oscillatory brain activity at the theta frequency in the rat hippocampus [97-98], suggesting that our findings can also be relevant to relaxin-mediated signaling in other organisms and systems.

#### PIL/GCL neurons as a new model to study the role of Cyclin A in postmitotic neurons

We find that PIL/GCL neurons stain positive for Cyclin A (CycA), a key cell-cycle protein, and require it for a proper response to the Dilp8 tissue-stress signal. Exactly how CycA contributes to Lgr3 function in PIL/GCL neurons remains to be defined. Nevertheless, our findings add to the increasing roles of cyclins in postmitotic cells. CycA has previously been implicated in regulating sleep amount together with other regulatory proteins, Rca1 and TARA, all in close connection with—but independently of— circadian pacemaker cells, providing a first hint towards a possible rhythm-control pathway [27]. The fact that both the adult CycA-positive neurons and the larval PIL/GCL-CycA positive neurons occupy similar brain areas (*pars lateralis* and *pars intercerebralis*) [20,27,28], which are homologous regions to the mammalian hypothalamus, support the relevance of this hypothesis. However, the requirement of Rca1 for sleep regulation, but not for Lgr3 activity in PIL/GCL neurons, suggests that CycA might regulate different processes in both situations.

A similar scenario is found in another non-replicative function of cyclins in postmitotic cells in vertebrates, where canonical and non-canonical cyclins, such as Cyclins A1, D1, and O, form mitotic oscillator-complexes together with their CDK catalytic subunits to participate in centriole amplification and ciliogenesis in multiciliated cells, independently of DNA replication [69-77]. In this pathway, whereas Emi1 (the vertebrate Rca1 ortholog), is dispensable [76], similarly to PIL/GCL neurons, another Emi1-paralogue assumes its role, so this is unlikely to represent a similar pathway. Moreover, in contrast to vertebrates, *Drosophila* cilia have only been documented in type I sensory neurons and sperm cells (as flagella) [99-102], so that CNS neurons–such as PIL/GCL neurons–are generally thought not to require centrosomal or ciliary proteins for their development and function [99-102]. It is important to consider, however, that the function of centrioles, centrosomes, and ciliary/basal body proteins have not yet been thoroughly assayed in specific CNS neurons with a defined developmental and physiological function in *Drosophila*, such as PIL/GCL neurons.

A more reasonable function for CycA in PIL/GCL neurons would be to regulate the activity of the postmitotic neuron CDK, CDK5, whose function has been extensively studied in postmitotic neurons, being involved in diverse processes from synaptic plasticity to circadian rhythm control (reviewed in Ref.[71]). Whereas, to our knowledge, CycA has never been found associated with CDK5 or other paralogues, if one considers the promiscuity of some cyclin-CDK interactions, PIL/GCL neuron CycA could assume similar roles as those proposed for Cyclin E and Cyclin Y, which regulate synaptic plasticity and axonal transport via the modulation of the activity of CDK5 or CDK5-like proteins in vertebrate and nematode postmitotic neurons, respectively [69-70]. Hence, further work directed to dissecting the role of CycA and how it relates to the intrinsic neuronal properties of the PIL/GCL neurons could possibly shed light not only into the mechanisms of the oscillatory response of PIL/GCL neurons to the Dilp8-dependent Lgr3 activation, but also to previously-reported postmitotic properties of cyclins, or novel, unanticipated roles for cyclins in postmitotic neurons.

As discussed above, a limitation of our study is that we cannot formally exclude a DNA replication role for CycA in PIL/GCL neurons. In *Drosophila* and vertebrates, CycA overexpression can result in ectopic S phases in different cell types [103-105], including a subset of postmitotic neurons of the developing eye [106]. If one considers that CycA levels are at least as high in PIL/GCL neurons as they are in other cycling cells, such as neuroblasts (**Figure 6B and Supplementary Figure S3**), one might hypothesize that PIL/GCL neurons could undergo DNA synthesis, or endocycles, and that this would be required for a proper response to Dilp8/Lgr3 activity. The major problem with this scenario is that endoreduplicating neurons have not been described in the larval CNS [78]. Nevertheless, endoreduplication is a well-described mechanism of cell growth in animals, including in neurons of other species [78]. The hypothesis here would be that the Dilp8/Lgr3 pathway either induces or requires CycA-dependent endoreduplication-related growth of the PIL/GCL neurons for proper activity. A second problem with this hypothesis is that it relies on extrapolating results from overexpression studies to the PIL/GCL neuron paradigm presented herein, which is nevertheless dependent on endogenous levels of CycA, not its ectopic or over-expression. In mitotically-dividing cells, such as *Drosophila* S2 cells in culture or in imaginal disc cells and stage 1-6 ovarian follicle cells *in vivo*, knocking-down CycA is sufficient to induce endocycles [107-110]. Such a function for CycA (to promote mitosis) in PIL/GCL neurons, would be paradoxal, as these are clearly postmitotic primary neurons, so we do not favor it. It is clear that the roles assumed by cyclins can be cell-type, context-(*i.e.,* mitotic or postmitotic), and species-specific. The strength of our findings, notwithstanding, is that we observe endogenous levels of CycA and perform loss-of-function experiments, which together suggest an endogenous role for CycA in postmitotic PIL/GCL neurons.

The new model that arises from our work (**Figure 8**), where Dilp8 induces PIL/GCL neuron oscillatory responses, and that these are critical for the behavioral change (delay in pupariation behavior) that promotes developmental stability, is both consistent with the intrinsic and neuroanatomical properties of PIL/GCL neurons and the physiological expectations of neurons with such properties.

**Figure 8.**
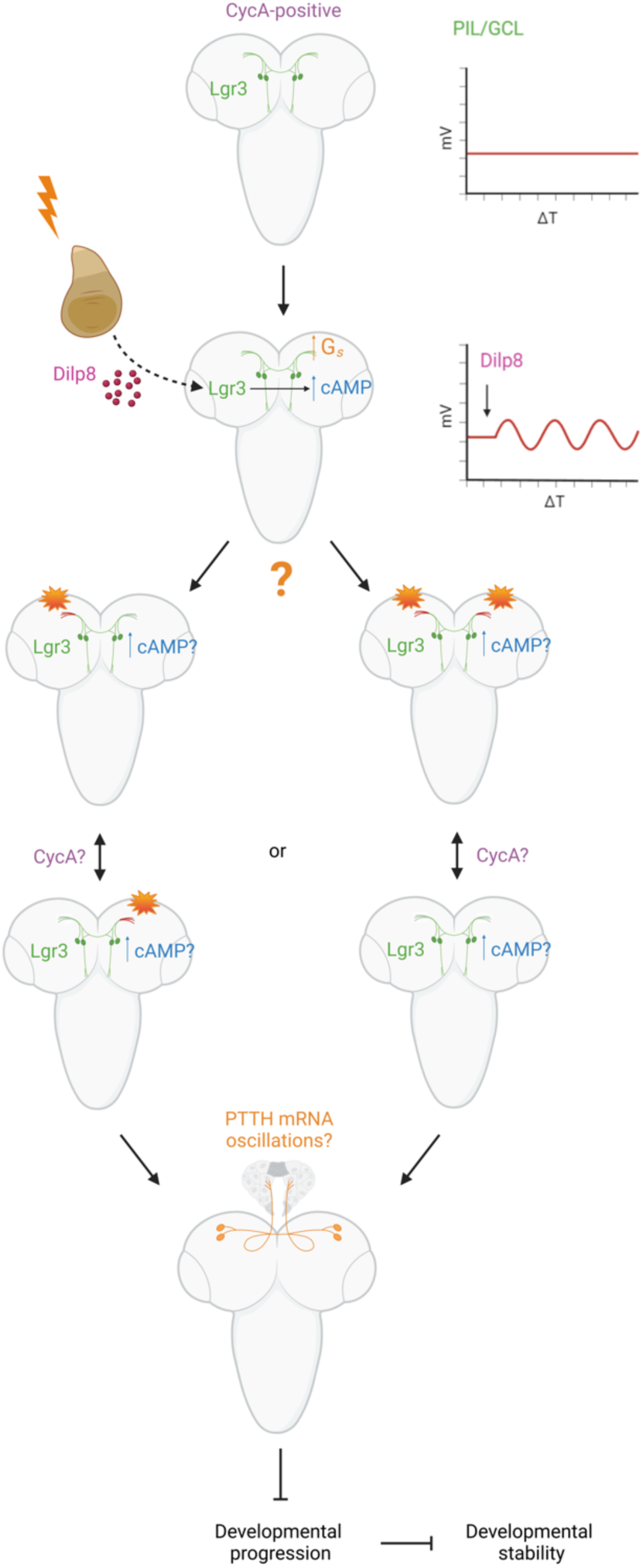
A new model for the role of PIL/GCL neurons in the Dilp8-Lgr3 developmental stability pathway. Upon imaginal disc damage (orange ray), imaginal disc cell abnormal growth triggers Dilp8 production and secretion (dark magenta circles), which activates Lgr3 in PIL/GCL neurons (green neurons). Lgr3 activity requires stimulatory G protein (G_s_, orange), which leads to an increase in cAMP levels (blue), and both depolarized and hyperpolarized states (oscillations in membrane potential, red curves in graphs, and orange flairs in the PIL/GCL neuron axons in the CNS schemes), which are required for the inhibition of the larval-to-pupal transition, which promotes developmental stability. In this model, it is unclear whether the oscillations occur contralaterally (left branch of the model) or bilaterally (right branch of the model), as well as their period. It also remains to be determined if oscillations arise from intrinsic pacemaker-like activity of PIL/GCL triggered by the Dilp8-Lgr3-G_s_-cAMP pathway (namely, that PIL/GCL neurons would oscillate even in isolation upon Lgr3 activation), or whether activation of this pathway leads to oscillations due to circuit architecture properties (feedback loops), which would mean that PIL/GCL neurons would only oscillate within their circuit context, not in isolation. The neuroanatomy of PIL/GCL neurons, a couple of bilateral local and projection neurons with a potential to form reciprocal pair loops, leads us to favor the latter model (right). The Dilp8/Lgr3 pathway and PIL/GCL neurons are proposed to act upstream of PTTH neurons (not experimentally addressed in this manuscript), which show oscillating production of *PTTH* mRNA in early 3rd instar larval periods. A tempting hypothesis is that endogenous Dilp8-Lgr3 promoted PIL/GCL neuron oscillations partially contribute to such *PTTH* mRNA oscillations during normal development and, in the presence of abnormal imaginal disc growth, the persistent Dilp8-Lgr3 signaling perpetuates this early-stage oscillatory pattern via prolonged PIL/GCL neuron oscillation, delaying the larval-to-pupal transition and promoting developmental stability.

#### Implication of the model to the Dilp8-Lgr3 developmental stability pathway. PIL/GCL neurons as regulators of downstream oscillations?

Here, we have tested widely accepted model of the Dilp8-Lgr3 pathway, confirming the initial predictions of the model, which had not yet been subject to specific genetic testing. Specifically, we have confirmed the role of *Lgr3* in PIL/GCL neurons, and the role of stimulatory G_s_ in the Dilp8-Lgr3 pathway, even though we could not show this latter requirement with the same specificity as we achieved for *Lgr3* due to technical reasons. These results are consistent with Lgr3 using Galphas signaling to increase cAMP levels via adenyl cyclase, and explains previous results using CRE-reporters and confirms the assumptions therein [20-21]. A connection between Galphas signaling, cAMP increase, and downstream activities of the PIL/GCL neurons remains to be made.

We then made the surprising observation that both depolarized and hyperpolarized PIL/GCL neuron states are required for their proper response to ectopic Dilp8 activity. This led to the proposition of a new model where PIL/GCL neurons respond to Dilp8 by becoming oscillators or becoming part of an oscillating circuit (**Figure 8**). The next neurons in this developmental stability neuroendocrine circuit are thought to be the PTTH neurons, with which PIL/GCL neurons make close contacts [19,21]. Larval PTTH neurons integrate many developmental and environmental signals, receiving input from different neuronal pathways, so that their regulation is complex [111-113]. PTTH neurons have the important function of producing and secreting PTTH [and other factors, such as the The Alk ligand Jelly belly (Jeb)], onto the prothoracic gland, which together contribute to the timely triggering of ecdysone biosynthesis [111,113-115]. The function of PIL/GCL neurons—when activated by Dilp8-Lgr3— would be to inhibit PTTH neuron function, thereby delaying the biosynthesis of ecdysone and consequently inhibiting developmental timing progression and the growth of imaginal discs in parallel [1-4,11]. Many questions arise from our work. For instance, how does Lgr3-Galphas-cAMP-signaling in PIL/GCL neurons mediate signaling to the PTTH-producing neurons, especially in the context of PIL/GCL neuron oscillation (**Figure 8**)? What would be the impact of such oscillations on PTTH neuron function?

Curiously, *PTTH* mRNA transcription shows an unusual cyclic pattern with an ∼8-h periodicity throughout the larval growth stage before the mRNA levels increase several fold as the larva approaches pupariation time [114]. Hence, oscillatory *PTTH* mRNA production is a characteristic of the larval growth stage. *PTTH* mRNA levels and their periodicity are controlled by the circadian pathway during this stage, albeit only partially [114]. Namely, *PTTH* mRNA production still oscillates with an increased period in *Pdf* mutants, suggesting that other mechanisms contribute to this oscillation. As the Dilp8-Lgr3 signaling pathway and PIL/GCL neurons are considered to act upstream of PTTH neurons and *PTTH* mRNA production in the developmental stability pathway [8-9,19,21,116], it is tempting to speculate that PIL/GCL neurons are instructive to this *Pdf-*independent mRNA oscillation, promoting and prolonging the early 3rd instar larval stage-like oscillations, thus counteracting the surge of *PTTH* at the end of the larval phase. In support for this hypothesis is the fact that endogenous *dilp8* mRNA production fluctuates during the 3rd instar larval stage in the absence of abnormal growth [9]. *dilp8* mRNA levels are over an order of magnitude lower under normal conditions than when there is imaginal disc abnormal growth (*e.g.,* [8-9]), but the mRNA levels are higher in earlier 3rd instar larvae than in latter stages [9]. This early Dilp8 production could theoretically lead to a developmentally-triggered Lgr3 activation in PIL/GCL neurons in early 3rd instar larvae, which could trigger PIL/GCL neuron oscillations, and hence contribute to the generation of the characteristic low level *PTTH* mRNA oscillation of earlier stages. This would also be consistent with previously proposed endogenous functions of the Dilp8-Lgr3 pathway, as revealed by anticipations of pupariation time in *dilp8* and *Lgr3* mutants in the absence of any genetic or environmental challenges, which implies that *PTTH* surges occurred precociously in these mutants [8-9,20].

## MATERIALS AND METHODS

### *Drosophila* husbandry and stocks

The stocks were reared in standard *Drosophila* food, and kept at 18 °C, ∼60%RH, 12-h light/dark cycle. All experiments were performed at 25 °C, 12-h light/dark cycle.

*sfGFP::Lgr3^ag^*^5^ was previously described [20]. *tsh-Gal80/CyO* (*r*ef.[41]) was a gift from Carlos Ribeiro. *w[1118]* and *Tub-dilp8(2)* and *(3)*, corresponding to 2nd and 3rd Chr insertions, respectively (ref.[21]), were gifts from Maria Dominguez. *MZ699-GAL4* (ref.[24]), *UAS-FRT>CD2>FRT-Kir2.1-GFP* (*UAS-FRT-stop-FRT-Kir2.1::GFP*) (ref.[47]), BL28842 w[*]; P{UAS(FRT.w[+mW.hs])TeTxLC}10/CyO (*UAS-FRT-stop-FRT-TeTxLC*) (ref.[50]), *BL28843* w[*]; P{UAS(FRT.w[+mW.hs])TeTxLC.IMPTNT}14A *UAS-FRT-stop-FRT-TeTxLC.IMPTNT*) (ref.[50]), and *w; UAS>stop>TrpA1/CyO* (*UAS-FRT-stop-FRT-TrpA1*) (ref.[51]) were gifts from Maria Luisa Vasconcelos and Marta Aranha. *UAS-TRPA1/CyO* was a gift from Christen Mirth. *w;;GMR19B09.1.1-GAL4, w;;GMR19B09.2.1-GAL4, w;;GMR19B09.3.1-GAL4, w;;GMR19B09.3A.1-GAL4, w;;GMR19B09.3B.1-GAL4, w;;GMR19B09.3C.1-GAL4, w;;GMR19B09.3D.1-GAL4,* and *w;;GMR19B09.3E.1-GAL4* (ref.[31]) were gifts from Geoffrey Meissner and Bruce Baker.

The following lines were obtained from the Bloomington Drosophila Stock Center at Indiana University:

BL36887 *y[1] sc[*] v[1] sev[21]; P{y[+t7.7] v[+t1.8]=TRiP.GL01056}attP2/TM3, Sb[1] (UAS-Lgr3-IR-V22)* (ref.[33])

BL48840 *w[1118]; P{y[+t7.7] w[+mC]=GMR19B09-GAL4}attP2* (ref.[31])

BL32222 *w[*]; P{y[+t7.7] w[+mC]=10XUAS-IVS-myr::tdTomato}attP40* (ref.[34])

BL32219 *w*; P{10XUAS-IVS-mCD8::RFP}attP40* (ref.[34])

BL55810 *w[1118]; P{y[+t7.7] w[+mC]=10XUAS(FRT.stop)GFP.Myr}su(Hw)attP5* (*UAS-FRT-stop-FRT-GFP*) (ref.[41])

BL38879 *P{w[+mC]=alphaTub84B(FRT.GAL80)}1, w[*]; Bl[1]/CyO; TM2/TM6B, Tb[1]* (ref.[43])

BL55819 *w[1118]; P{y[+t7.7] w[+mC]=8XLexAop2-FLPL}attP2* (ref.[34])

BL66675 *w[*]; P{w[+mC]=UAS(FRT.stop)shi[ts]}2* (ref.[50])

BL66676 *w[*]; P{w[+mC]=UAS(FRT.stop)shi[ts]}3* (ref.[50])

BL33064 *w1118; P{UAS-DenMark}2, P{UAS-syt.eGFP}2; In(3L)D, mirrSaiD1 D1/TM6C, Sb1* (ref.[75])

BL64089 *w[1118] P{y[+t7.7] w[+mC]=R57C10-FLPG5.PEST}su(Hw)attP8; PBac{y[+mDint2] w[+mC]=10xUAS(FRT.stop)myr::smGdP-HA}VK00005 P{y[+t7.7] w[+mC]=10xUAS(FRT.stop)myr::smGdP-V5-THS-10xUAS(FRT.stop)myr::smGdP-FLAG}su(Hw)attP1* (ref.[79])

BL55752 *w[*]; P{y[+t7.7] w[+mC]=LexAop-tdTomato.Myr}su(Hw)attP5; P{w[+mC]=UAS-FLP.D}JD2, PBac{y[+mDint2] w[+mC]=brp(FRT.Stop)V5-2A-LexA-VP16}VK00033* (ref.[80])

BL682125 *w[*]; P{w[+mC]=Pdfr-GAL4.B}2/CyO* (ref.[63])

BL7127 *y[1] w[*]; P{w[+mC]=GAL4-per.BS}3* (ref.[60])

BL50704 *y[1] v[1]; P{y[+t7.7] v[+t1.8]=TRiP.HMC03106}attP2* (*UAS-Galphas-IR*) (ref.[33])

BL31761 *y[1] v[1]; P{y[+t7.7] v[+t1.8]=TRiP.HM04073}attP2 (UAS-CycA-IR)* (ref.[33])

BL51450 *y[1] sc[*] v[1] sev[21]; P{y[+t7.7] v[+t1.8]=TRiP.HMC03180}attP2* (*UAS-Rca1-IR*) (ref.[33])

### Immunofluorescence analyses

CNS of larvae L3 wandering were dissected in Schneider Medium (Biowest – cat. #L0207 or Gibco - cat. #21720-024), fixed for 30 min in 4% paraformaldehyde, rinsed with PBS with Triton (0.3%) (PBST), incubated with primary antibody for 24-48h and with fluorescently labeled secondary antibody for 2–24 h in PBST with 1% bovine serum albumin. Samples were washed 3x for 30 min each in PBST after each antibody incubation. Nuclei were counterstained with DAPI (Sigma) and tissues were mounted in DABCO mounting medium. Primary antibodies used were: Rabbit Anti-GFP, 1:200 (Life Technologies, A11122); Rabbit Living Colors DsRed, 1:500 (Clontech, 632496); Mouse Anti-GFP-4C9, 1:200 (DSHB, DSHB-GFP-4C9), Mouse nc82-s, 1:100 (DSHB, AB_2314866), Mouse anti Cyclin A (A12), 1:100, (DHSB, AB_528188), Rat DN-Ex #8-st, 1:50 (DSHB, AB528121). Secondary antibodies were: Alexa Fluor 488, anti-mouse (Life Technologies, A11029); Alexa Fluor 488, anti-rabbit (Life Technologies, A11070); Alexa Fluor 488, anti-rat (Invitrogen, A21208), Alexa Fluor 568, anti-rabbit (Life Technologies, A11036); Alexa Fluor 594, anti-mouse (Life Technologies, A11020), Alexa Fluor 647 anti-rabbit (Jackson Immuno Research, 711-605-152). Images were obtained with a Zeiss LSM 710 or a Leica SP5 Confocal Microscopes. Images were analyzed using FIJI software [117]. Usually, 5 to 10 CNS were mounted for observation and 1 representative image per genotype is depicted in figures. CNS from male and female larvae were scored together.

### Measurement of the developmental timing of pupariation

Male and virgin female flies aged 3-15 d were crossed and 48 h later transferred to an agar plate with yeast paste. The next morning, the flies were transferred to a fresh plate to lay eggs for 2 h. The plates were exchanged every 2 h, 4-5 times a day. Second instar larvae (48 h after egg laying) were transferred to vials containing standard *Drosophila* food. To prevent overcrowding, the number of larvae per vial was between 20 to 30. Survey of pupae consists in counting the number of puparia in each time interval. The final number of pupae for each genotype in each experiment is depicted in the respective figures. For pupariation time assays, we aimed at obtaining at least 30 individual larvae per genotype, yet some genotypes were sick or difficult to obtain due to balancer chromosomes, yielding fewer larvae of the correct genotype to score. Males and females were scored together. These experiments were performed at 25 ± 1 °C, except for experiments using flies carrying the *TrpA1* and *shi[ts]*, which were performed at 18 °C until the second instar (72 h after egg laying) after which the larvae were transferred to 29 °C to activate the system.

### General study design and statistics

In all experiments reported in this work, no data point was excluded. All data points, including outliers, are represented in the figures and were used in the statistical analyses. No blinding was done and no particular randomization method was used to attribute individuals to experimental groups. Statistics were performed in RStudio (RStudio: Integrated Development for R. RStudio, PBC, Boston, MA URL http://www.rstudio.com/). Graphs were made in GraphPad Prism version 8.0.2 for Windows, GraphPad Software, Boston, Massachusetts USA, www.graphpad.com. Scheme art in Figures 1-3, 5, 7, and 8 was partially done in Biorender.com and Inkscape.org v1.4.

## AUTHOR CONTRIBUTIONS

R.Z., J.P., M.F-A., M.P., L.L., A.P.C., A.G., and F.H. performed genetic, phenotypic, and behavioral experiments, and contributed to image and data analyses. R.Z., A.G., F.H., and A.M.G. designed research. A.M.G. contributed to image and data analyses. F.H. and A.M.G. supervised the work. A.M.G. wrote the manuscript with the help of all authors.

## ACKNOWLEDGEMENTS

We thank Drs. Maria Luisa Vasconcelos, Carlos Ribeiro, Christen Mirth, Maria Dominguez, Geoffrey Meissner, and Bruce Baker for fly stocks. We thank lab members, Dr. João Picão, Dr. Filipe de Sousa, Ana Sofia Abril, Sofia Pinto, Leonor Bezerra, and Tiago Andrade for discussions and/or comments on the manuscript. Stocks obtained from the Bloomington Drosophila Stock Center (NIH P40OD018537) were used in this study. TRiP RNAi stocks were used in this study (Office of the Director R24 OD030002). The monoclonal antibodies Anti-GFP-4C9, nc82-s, anti-Cyclin A(A12), and DN-Ex #8-st developed by DSHB, E. Buchner, Christian Lehner, and T. Uemura, respectively, were obtained from the Developmental Studies Hybridoma Bank, created by the NICHD of the NIH and maintained at The University of Iowa, Department of Biology, Iowa City, IA 52242. Work in the Integrative Biomedicine Laboratory was supported by the European Commission FP7 (PCIG13-GA-2013-618847), by the FCT (PTDC/BEXBCM/1370/2014; PTDC/BIA-BID/31071/2017; PTDC/MED-NEU/30753/2017 (LISBOA-01-0145-FEDER-030753, with financing from the Programa Operacional Lisboa 2020); EXPL/BIA-BID/1524/2021; EXPL/BIA-COM/1296/2021; 10.54499/2022.03859.PTDC; 2023.15344.PEX), by the UIDs iNOVA4Health (10.54499/UIDB/04462/2020) and cE3c (10.54499/UIDB/00329/2020), by LS4FUTURE (10.54499/LA/P/0087/2020) and CHANGE (10.54499/LA/P/0121/2020) financed by the FCT/MCTES (Portugal), and by Congento LISBOA-01-0145-FEDER-022170, co-financed by FCT/Lisboa2020; UID/Multi/04462/2019; and by the Microscopy Facility of FCUL (PPBI-POCI-01-0145-FEDER-022122). Work in the Garelli lab was also supported by ANPCyT (Agencia Nacional para la Promoción de la Ciencia y la Tecnología, PICT 2017-0254 and PICT 2020-01568), CONICET (PIP11220150100182CO) and Universidad Nacional del Sur PGI 24/B288. AMG and FH were individually supported by grants 10.54499/CEECINST/00102/2018/CP1567/CT0031 and 10.54499/DL57/2016/CP1457/CT0016, respectively. AG is a CONICET researcher.

