## Supplementary Material for "Neurogenetic evidence that Dilp8 promotes developmental-stability via Lgr3-neuron oscillation"

[1] cE3c - Centre for Ecology, Evolution and Environmental Changes & CHANGE - Global Change and Sustainability Institute, Department of Animal Biology, Faculty of Sciences, University of Lisbon, Lisbon, Portugal.
[2] iNOVA4Health, Nova Medical School, Faculdade de Ciências Médicas, NMS, FCM, Nova University of Lisbon, Lisbon, Portugal.
[3] Institute of Molecular Biology, Mainz, Germany.
[4] Laboratorio de Genética del Comportamiento, Fundación Instituto Leloir - Instituto de Investigaciones Bioquímicas de Buenos Aires (IIBBA- CONICET), Buenos Aires, Argentina.
[5] Instituto de Investigaciones Bioquímicas de Bahía Blanca (INIBIBB), CONICET and Departamento de Biología, Bioquímica y Farmacia, Universidad Nacional del Sur, Bahía Blanca, Argentina.
* These authors contributed equally to this work.

**This file contains:**
**Supplementary Figures S1-S6.**
**Supplementary References.**

#### Supplementary Figure S1

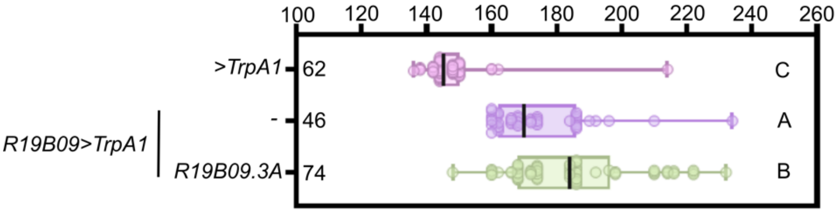


**Supplementary Figure S1. Thermogenetic activation of *R19B09.3A>* cells does not negatively affect the delay caused by thermogenetic activation of *R19B09>* cells.** Box plot showing pupariation time (Time after egg laying (AEL)) in h of (N) larvae expressing *TrpA1* under the control of *R19B09>* or *R19B09>* and *R19B09.3A>* together. Negative control (blue): *>TrpA1* crossed with *w[1118]*. Whiskers extend to minimum or maximum values. Each dot represents one individual. The vertical bars represent the median. $P < 0.0001$, One-way ANOVA test. Genotypes sharing the same letter are not statistically different at $\alpha = 0.05$, Tukey's *post-hoc* test.

#### Supplementary Figure S2

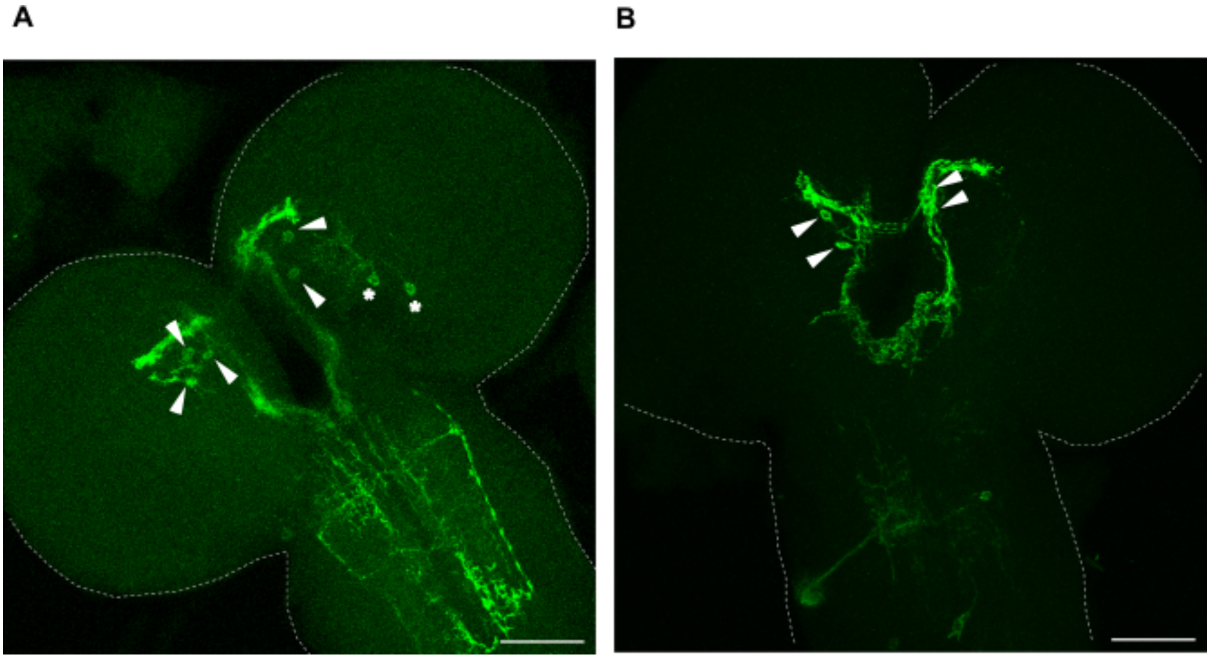


**Supplementary Figure S2. (A)** Examples of other brain neurons inconsistently labeled by the genetic intersection of *R19B09* and *MZ699* elements (*R19B09* ∩ *MZ699*) (asterisks). Cells in the brain region corresponding to PIL/GCL neurons (arrowheads). Shown is the max intensity projection of confocal z-stack slices of an immunohistochemistry preparation of dissected wandering L3 larval CNSs (anterior contour depicted with a dashed white line) expressing *R19B09* ∩ *MZ699>GFP::myr* stained with anti-GFP (green). **(B)** An apparent neuroblast lineage inconsistently labeled by *R19B09* ∩ *MZ699>*. Same conditions as (A). Scale bars, 50 µm.

#### Supplementary Figure S3

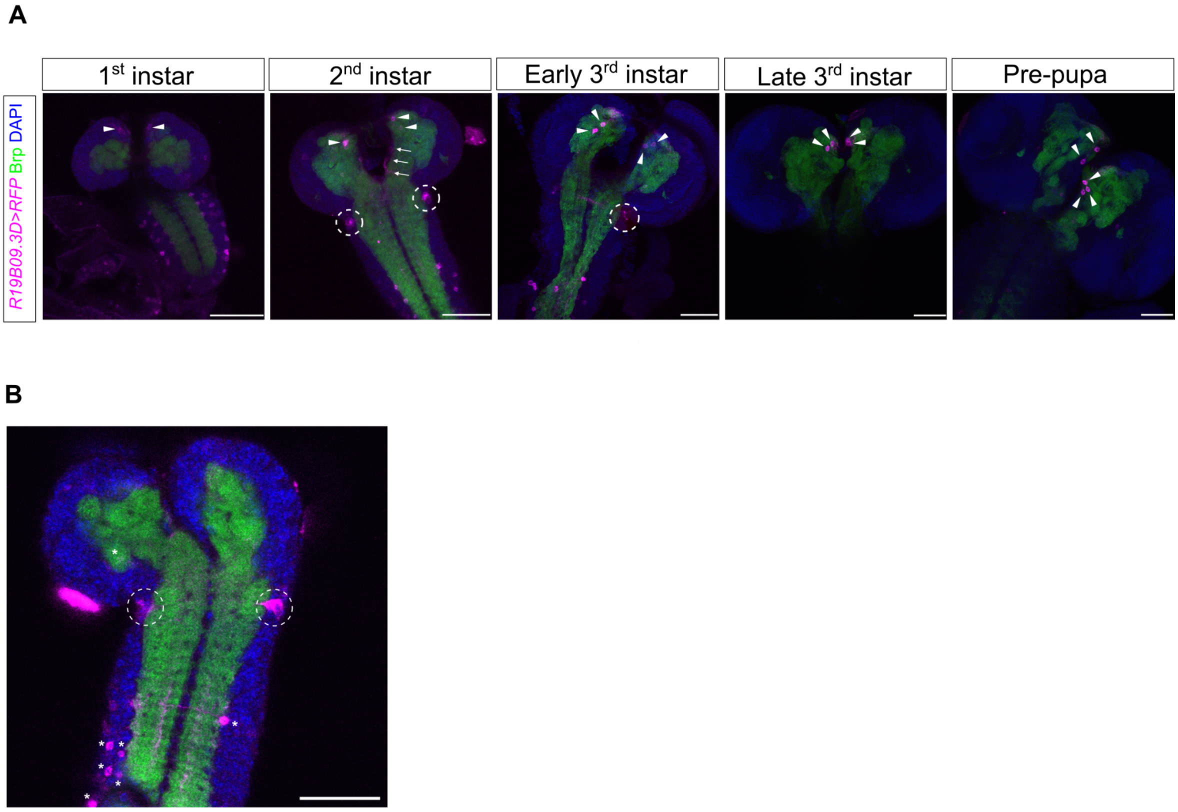


**Supplementary Figure S3. PIL/GCL neuron neuroanatomy during development. (A)** Sum of confocal z-stack slices of CNSs from different larval stages stained with nc82 (anti-Brp) (neuropil, green) and DAPI (nuclei, blue). The *Lgr3* CRM *R19B09.3D>* drives myr::RFP expression (*R19B09.3D>mCD8::RFP*, magenta). Arrowheads label PIL/GCL neurons. Arrows depict visible descending projections from the PIL/GCL region in 2nd instar larvae. The other projections are not visible in these confocal stack projections. Dotted circles highlight neuroblast lineages. **(B)** Possible neuroblast lineage labeled by *R19B09.3D>* (and *PIL/GCL>*) (dotted circles). Shown is a single ventral slice of an L2 larval stage CNS. Same conditions as in (A). VNC neurons, asterisks. Scale bars, 50 µm.

#### Supplementary Figure S4

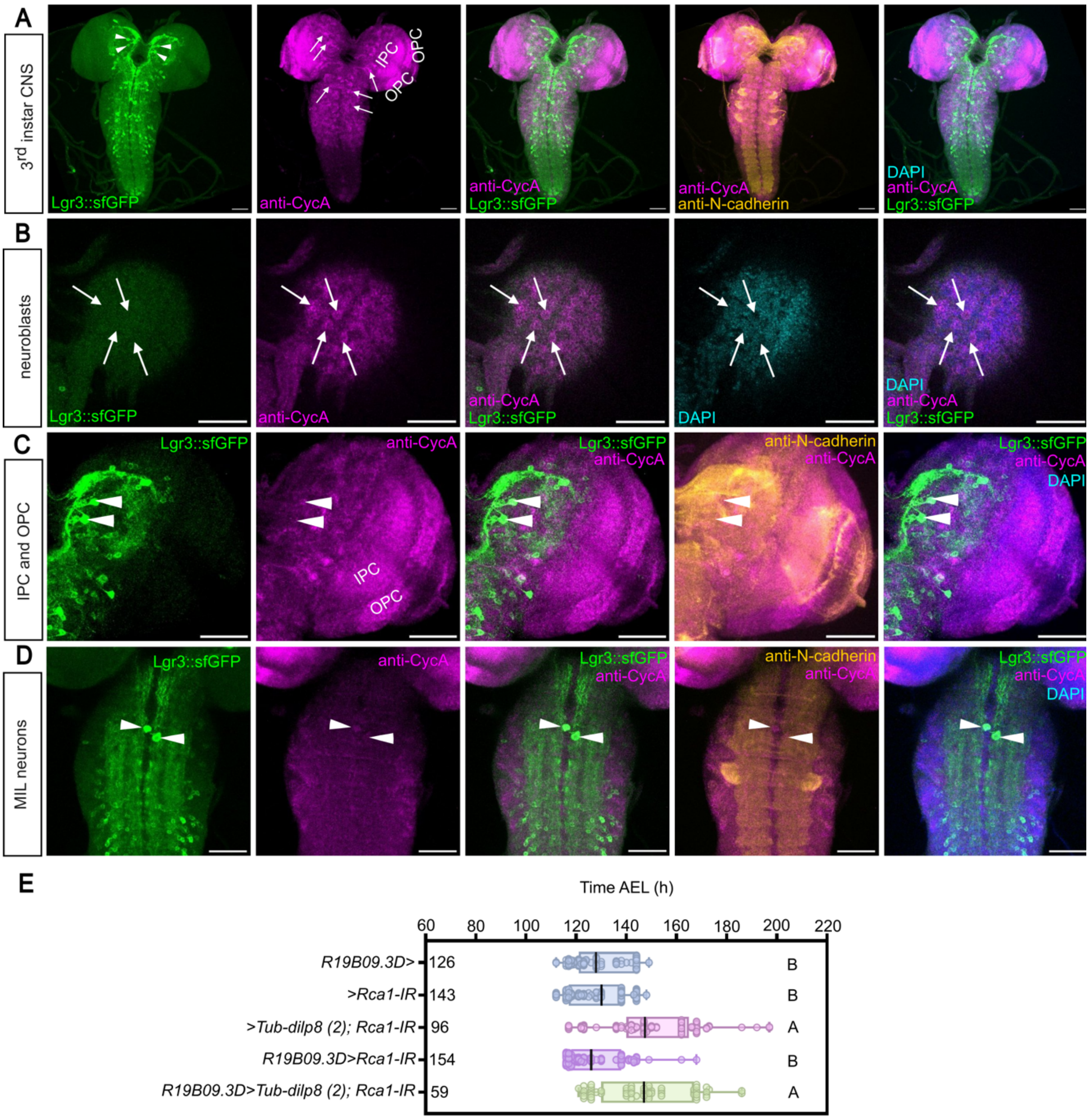


**Supplementary Figure S4. Neuroblasts, inner (IPC) and outer (OPC) proliferative centers, and Midline Internal Lgr3-positive (MIL) neurons stain positively for CycA. (A)** General pattern of CycA-positive cells in the 3rd instar larva CNS. Inner (IOC) and outer (OPC) proliferative centers and several neuroblast/lineages (arrows) are visible in the magenta channel (anti-CycA). Depicted is the sum of confocal z-stack slices of an immunohistochemistry preparation obtained from a dissected 3rd instar larva CNS stained with anti-GFP (sfGFP::Lgr3, green), anti-CycA (magenta), anti-DN-Ex#8-s (anti-N-cadherin) (neuroanatomical reference, yellow), and counterstained with DAPI (nuclei, blue). Arrowheads label PIL/GCL neurons. **(B)** Same as (A) yet with less slices per projection and focusing on the superficial optic lobe area to depict neuroblasts positively stained for CycA (arrowheads). **(C)** Same as (A), with a close-up of the right optic lobe area showing that PIL/GCL neurons (arrowheads, green), IPC, and OPCs stain positively for anti-CycA (magenta). Arrowheads label PIL/GCL neurons. **(D)** Same as (A), with a close-up of the thoracic region of the VNC showing that MIL neurons (arrowheads) stain positively for anti-CycA. These images are taken from the same CNS depicted in Figure 6. Scale bars, 50 µm. **(E)** Downregulation of the *Rca1* (*Rca1-IR*) in *R19B09.3D>* cells does not rescue the delay induced by *Tub-dilp8(2)*. Negative controls (blue), *R19B09.3D>* or *>Rca1-IR* crossed with *w[1118]*. This experiment was conducted in parallel with the ones reported in Figures 4F and 6E, so that the *R19B09.3D>* animals are the same in the three figures panels. $P < 0.001$, One-way ANOVA test. Genotypes sharing the same letter are not statistically different at $\alpha = 0.05$, Tukey's post-hoc test.

#### Supplementary Figure S5

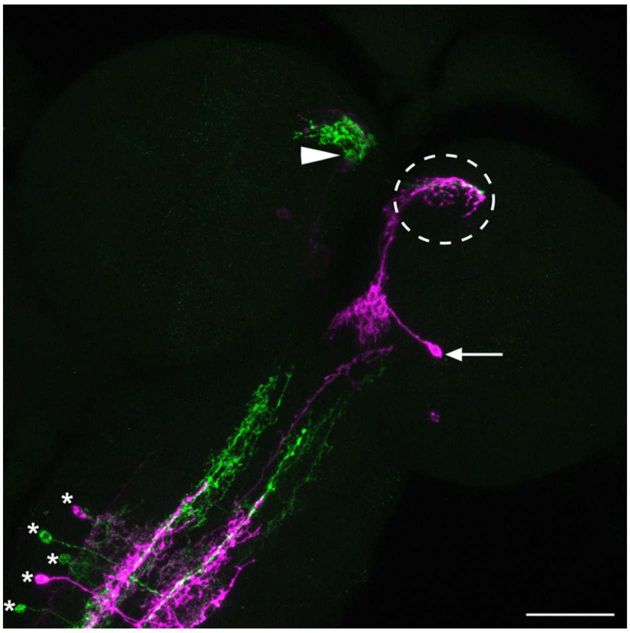

**Supplementary Figure S5. A frequently observed posterior brain neuron (magenta soma, arrow) in *R19B09.3D>* MCFO clones.** This neuron sends an anterior projection to the same compartment (magenta in dotted circle) receiving contralateral innervation from a PIL/GCL neuron in the other brain hemisphere (green). The PIL/GCL neuron soma is depicted with an arrowhead. Depicted is the sum of confocal z-stack slices of MCFO labeling of L3 larval stage neurons with panneuronally (*R57C10*>)-driven expression of *FLP* and *R19B09.3D*>-driven expression *UAS*-driven MCFO markers differentially labeled here with anti-GFP (green) and anti-V5 (magenta). Asterisks denote other VNC neurons. Scale bar, 50 µm.

#### Supplementary Figure S6

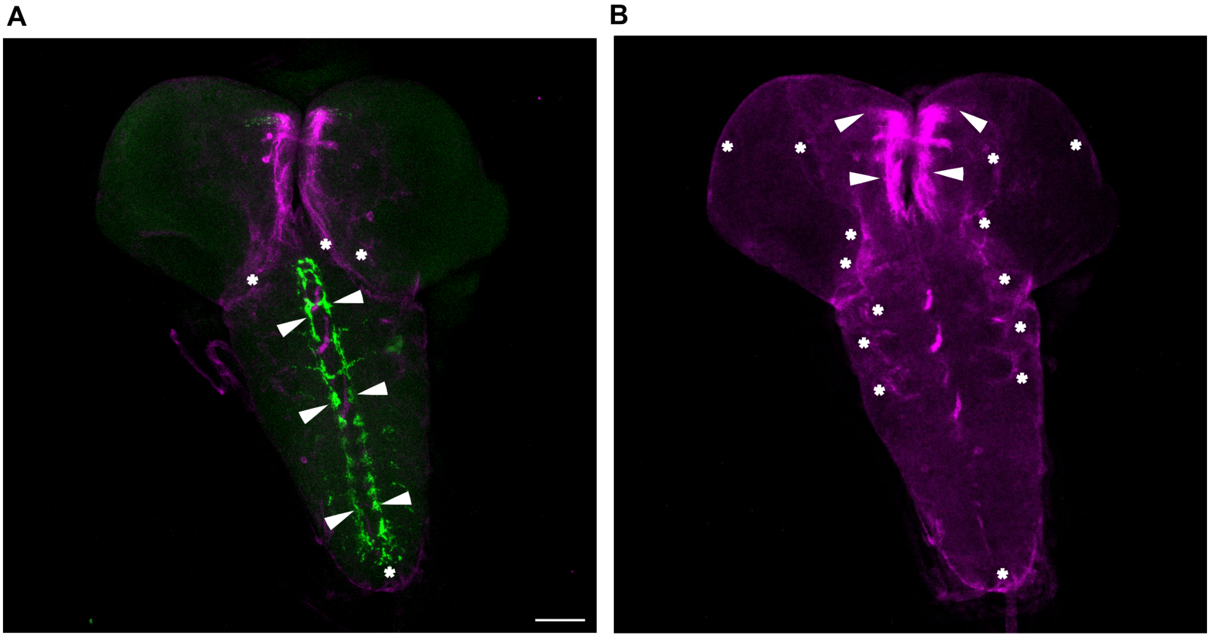


**Supplementary Figure S6. Full CNS confocal projection of the brain region highlighted in Figure 7E. (A)** Dendritic and axonal compartments of *PIL/GCL>* cells labeled with somatodendritic marker, DenMark (magenta), and the synaptic vesicle marker, syt::GFP (green), respectively. Apart from the features highlighted in **Figure 7E**, and the occasional cells already described in the text, this experiment revealed previously unnoticed cell types labeled by the *PIL/GCL>* driver. We noticed 1) diffuse DenMark staining in cells that occupy very large superficial areas, which could be a subset of glial cells (asterisks) and 2) syt::GFP labeling in axonal terminations in the VNC that likely correspond to axonal termini of Class IV multidendritic sensory neurons (arrowheads; Grueber *et al.,* Development, 2007). The reason why these went unnoticed are two-fold, first diffuse glial expression might be hard to see when other stronger neurons are labeled. Second, the sensory projections are only evident here because of the strong syt::GFP label and the removal of other syt::GFP expression from the VNC by the *tsh-GAL80* construct present in the *PIL/GCL>* driver. We believe that none of these cells are critical for the Lgr3-dependent effects we are studying because Lgr3 RNAi in glial cells using the pan-glial *repo-GAL4* driver does not rescue the Dilp8-dependent delay (Colombani *et al.,* Curr Biol, 2015), and other strongly suppressing drivers such as *R19B09.3D>* does not drive detectable expression in such Class IV sensory neurons (see for instance, **Supplementary Figure S3B**, where there are no such ladders or patterns in the VNC). **(B)** Max intensity projection of a subset of the slices of the *PIL/GCL>DenMark* channel to capture the glial-cell like pattern (asterisks) in addition to the soma of and descending dendrites from the PIL/GCL neuron area (arrowheads). Notice the superficial staining around the CNS, the neuropil, and the lateral sides of the leg neuropil compartments. Scale bar, 50 µm.
